# Cell cycle-driven epigenetic resetting maintains stem cell fate for shoot branching

**DOI:** 10.1101/2025.10.22.683900

**Authors:** Yahe Guo, Yi Yang, Bihai Shi, Hongli Wang, Xiuwei Cao, Ziyuan Peng, Yuehui He, Jun Xiao, Louis Tao, Doris Wagner, Masaaki Umeda, Robert Sablowski, Ying Wang, Yuling Jiao

## Abstract

Adult mammalian stem cells typically maintain stem cell identity through proliferative quiescence. In contrast, we demonstrate that stem cell maintenance in *Arabidopsis* bud precursor cells requires active cell-cycle progression. Inhibiting division silences the shoot meristem marker gene *SHOOT MERISTEMLESS* (*STM*) and promotes differentiation. Whereas proliferation dilutes H3K27me3 levels to counteract silencing. Meanwhile, we identified two classes of transcription factors recruiting polycomb repressive complex 2 (PRC2) to epigenetically silence *STM*. The balance between these forces establishes a cell cycle-coupled epigenetic “Sisyphus” mechanism that maintains pluripotency. This cell fate switch is bistable; modeling and experimental data confirm that prolonged quiescence triggers irreversible differentiation. We propose that sequence-dependent PRC2 recruitment in plants enables precise silencing of fate-determining genes, while cell proliferation sustains pluripotency by resetting epigenetic marks.

## Introduction

Adult stem cells maintain tissue function and integrity throughout an organism’s lifespan. In mammals, a hallmark of adult stem cells is their proliferative quiescence (*1*, *2*). This functionally critical state enables slower-cycling hematopoietic (*3*), neural (*4*), and intestinal (*5*) stem cells to achieve long-term maintenance, while their fast-cycling counterparts undergo rapid exhaustion or premature differentiation. In plants, adult stem cells play a similar role in maintaining tissue homeostasis and driving postembryonic organogenesis (*6*), including shoot branching. Axillary bud progenitor cells (AxPs) located in the leaf axil (*7*) sustain expression of the shoot meristem marker *SHOOT MERISTEMLESS* (*STM*) (*8*, *9*) and form axillary meristems (AMs) through cytokinin-activated *de novo WUSCHEL* (*WUS*) expression (*10*), ultimately achieving shoot apical meristem (SAM)-equivalent developmental potential (*11*). Although AxPs reside in slow-dividing boundary regions (*12*, *13*) and prolonged quiescence has been presumed essential for stem cell maintenance (*14*), we demonstrate that AxPs actually divide actively and that slowed proliferation instead accelerates differentiation. Proliferating AxPs maintain *STM* expression accompanied by reduced enrichment of histone H3 with Lys27 trimethylation (H3K27me3). We further identify transcription factors that recruit polycomb repressive complex 2 (PRC2) to deposit H3K27me3 at the *STM* locus. Crucially, H3K27me3 dilution during cell division establishes a bistable epigenetic switch: only dividing AxPs sustain low H3K27me3 levels and *STM* expression, thereby constituting a cell cycle-coupled epigenetic “Sisyphus” mechanism.

## Results

### AxPs maintain slow cell proliferation

AxPs reside in the center of the boundary region within the leaf axil and exhibit minimal rates of cell division (*13*). To precisely monitor their cell cycle progression, we fluorescently labeled dividing cells via an EdU assay (*15*). Active DNA synthesis was prominently observed in the SAM and leaf primordia, but was sparse in the leaf axils (Fig. 1A). Crucially, S-phase progression persisted, especially in leaf axil centers (Fig. 1, B and C).

**Fig. 1.**
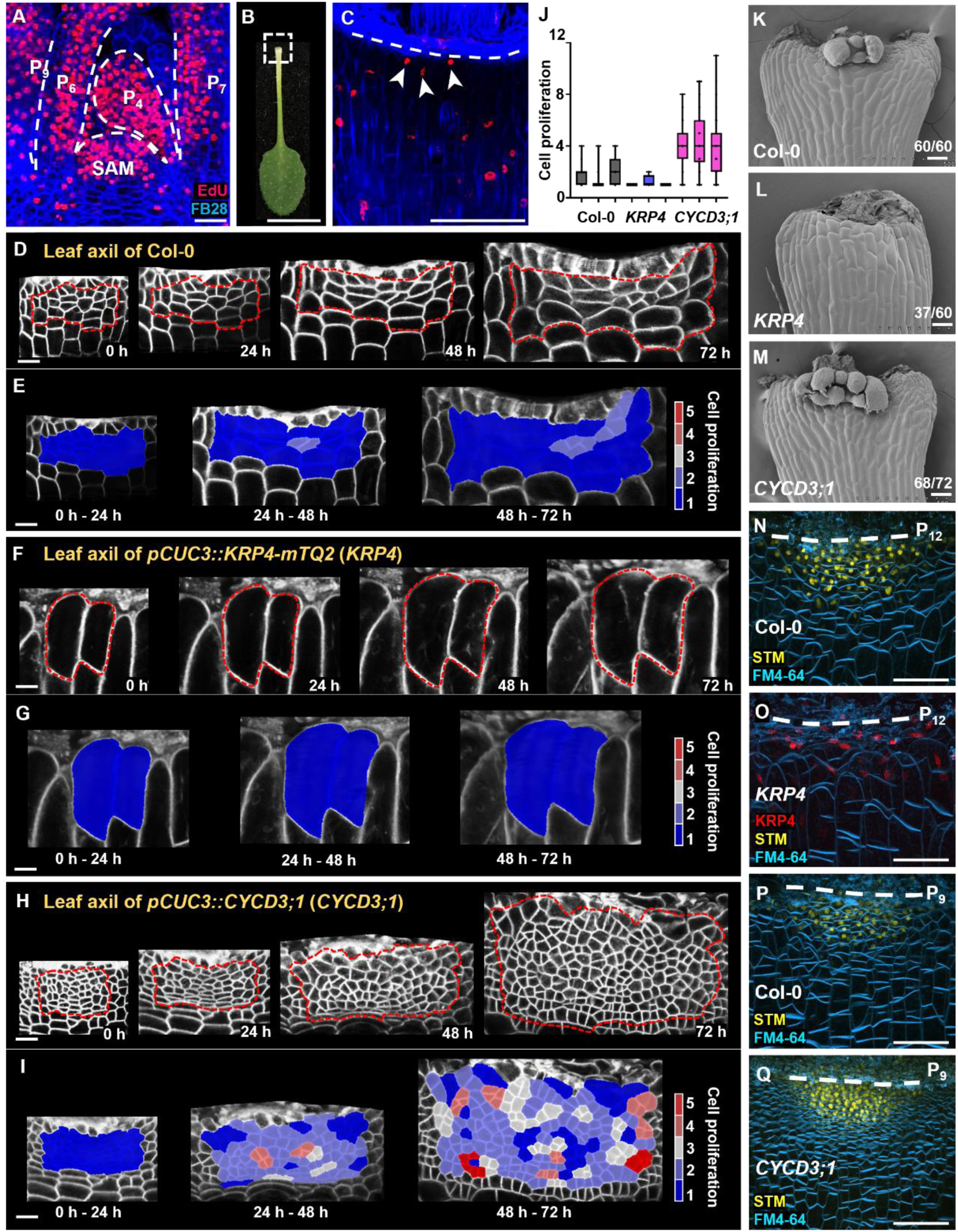
Slow cell division is maintained in the leaf axils. (**A**) Longitudinal sections of the vegetative shoot apices of Col-0 plants showing the EdU-stained cell nuclei (red) and FB28-stained cell walls (blue). The white dashed lines delineate the boundary area between the two leaf primordia. Note the existence of EdU signals in the leaf axil. Scale bars, 100 µm. (**B** and **C**) Live imaging showing cell division at the leaf axils of P_9_. The white dotted box in (B) indicates the leaf axils imaged in (C). EdU-stained cell nuclei (red), and FB28-stained cell walls (blue). The white dashed line in (C) marks the incision line. The white arrows in (C) indicate dividing nuclei. Scale bars, 1 cm in (B), and 100 µm in (C). (**D, F, H**) Time-lapse images of detached P_9_ leaf axil cells from Col-0 (D), *pCUC3::KRP4*-*mTQ2* (*KRP4*, F) and *pCUC3::CYCD3;1* (*CYCD3;1*, H) plants. The L1 layer of the leaf axil from short-day-grown plants was reconstructed using MorphoGraphX. The red dashed boxes highlight the leaf axil cells used for lineage tracking, with an approximate area of 3000 µm^2^. Scale bars, 20 µm. (**E, G, I**) Heatmaps showing the cell division rates in the red dashed boxes in (D), (F) and (H), respectively. Scale bars, 20 µm. (**J**) Statistical analysis of cell division within the red dashed boxes over 72 h showing differences among Col-0, *KRP4* and *CYCD3;1*, each with three biological replicates. (**K**, **L, M**) Axillary bud initiation in detached Col-0 (K), *KRP4* (L) and *CYCD3;1* (M) leaves. Note the lack of axillary buds in *KRP4* and the formation of two axillary buds in *CYCD3;1*. (m/n) indicates that m in n biological repeats shows the displayed features. Scale bars, 100 µm. (**N** and **O**) Leaf axil *STM* signals (yellow) in Col-0 (N) and *pCUC3::KRP4*-*mTQ2* (*KRP4*, O) in P_12_, with FM4-64 staining (cyan). Red indicates *KRP4* expression (O). Note the lack of *STM* signals in *KRP4*. Scale bars, 50 µm. (**P** and **Q**) Leaf axil *STM* signals (yellow) in Col-0 (P) and *CYCD3;1* (Q) in P_9_, with FM4-64 staining (cyan). Note the enhanced *STM* signals in *CYCD3;1*. Scale bars, 50 µm.

Spatiotemporal analysis of cell division in the leaf axil, conducted through time-lapse imaging, revealed that AxPs undergo division, in contrast to the quiescent flanking boundary cells. At the bases of the ninth youngest leaf primordium (P_9_), where AMs have not yet formed, we conducted live imaging of AM initiation and bulging over four days (Fig. 1D). Following cell segmentation and lineage tracing, growth properties were quantified (Fig. 1E and fig. S1A). Initially, cell division and expansion proceeded slowly, but then transitioned to accelerated AxP proliferation accompanied by cell expansion just prior to AM formation (Fig. 1, D and E, and fig. S1A). This indicates that AxPs sustain low cell proliferation rates until AM initiation, at which point division accelerates.

### AM initiation requires cell division

To investigate how cell proliferation influences stem cell fate, we generated transgenic lines with altered boundary cell division rates. Precise inhibition of division was achieved by expressing the G1/S transition inhibitor KRP4/ICK7(*16*, *17*) fused to mTurquoise2 (mTQ2) under the boundary-specific *CUP-SHAPED COTYLEDON3* (*CUC3*) promoter (*18*). In *pCUC3::KRP4-mTQ2* leaf axils, cell division was strongly suppressed with rare mitotic events (Fig. 1, F, G and J), yielding enlarged cells (fig. S1B). Unlike in the wild type, most mature *pCUC3::KRP4-mTQ2* leaf axils lacked axillary buds (fig. S1E), suggesting AxP maintenance defects. When cultured, detached wild-type leaves reproducibly formed axillary buds or branches within 3 d, while *pCUC3::KRP4-mTQ2* leaves rarely produced any buds (Fig. 1, K and L, and fig. S1D).

Conversely, expression of the G1/S-promoting cyclin *CYCD3;1* (*16*, *19*) driven by the *CUC3* promoter accelerated cell division in the leaf axil and led to earlier AM formation (Fig. 1, H, I and J), accompanied by a reduction in cell size (fig. S1C). The rapid proliferation and expansion of the area prevented long-term lineage tracking. In contrast to the effects of *KRP4*, *pCUC3::CYCD3;1* plants developed additional accessory buds (fig. S1E). In cultured detached leaves, which already show slightly accelerated bud formation (*9*), individual *pCUC3::CYCD3;1* axils frequently formed 3 to 5 buds (Fig. 1, K and M, and fig. S1D). Enhanced AM initiation was further confirmed in *krp3467* quadruple mutants, which exhibited increased accessory bud formation (fig. S1, F to K). Together, these results demonstrate that leaf axil cell proliferation is essential for AM initiation and that accelerating proliferation promotes ectopic AM initiation.

### STM maintenance in AxP depends on cell cycle progression

AM initiation requires sustained *STM* expression within AxPs (*8*, *9*). When assessing cell fate in *pCUC3::KRP4-mTQ2* plants, we observed significantly weakened and faster-diminishing *STM* signals (fig. S2, A and B). The majority of *pCUC3::KRP4-mTQ2* leaf axils at the P_12_ stage lacked detectable *STM* expression (Fig. 1, N and O). Furthermore, while nuclei in wild-type P_15_ leaf axils maintained a rounded morphology, the nuclei in *pCUC3::KRP4-mTQ2* cells adopted an elongated, fusiform shape (fig. S2, D and E), a characteristic indicative of cellular differentiation (*20*).

Analysis of key regulators for AM initiation showed complete absence of expression for both *REGULATOR OF AXILLARY MERISTEMS1* (*RAX1*) and *REVOLUTA* (*REV*) in *pCUC3::KRP4-mTQ2* P_12_ leaf axils (fig. S3, A, B, D and E), both of which are essential for AM initiation (*21–23*). Additionally, *WUS*, which is activated *de novo* just prior to AM initiation (*10*), was also undetectable in these plants (fig. S3, C and F), consistent with their failure to form axillary buds. These findings collectively indicate that suppressing cell division severely impairs the maintenance of stem cell identity in AxPs.

**Fig. 2.**
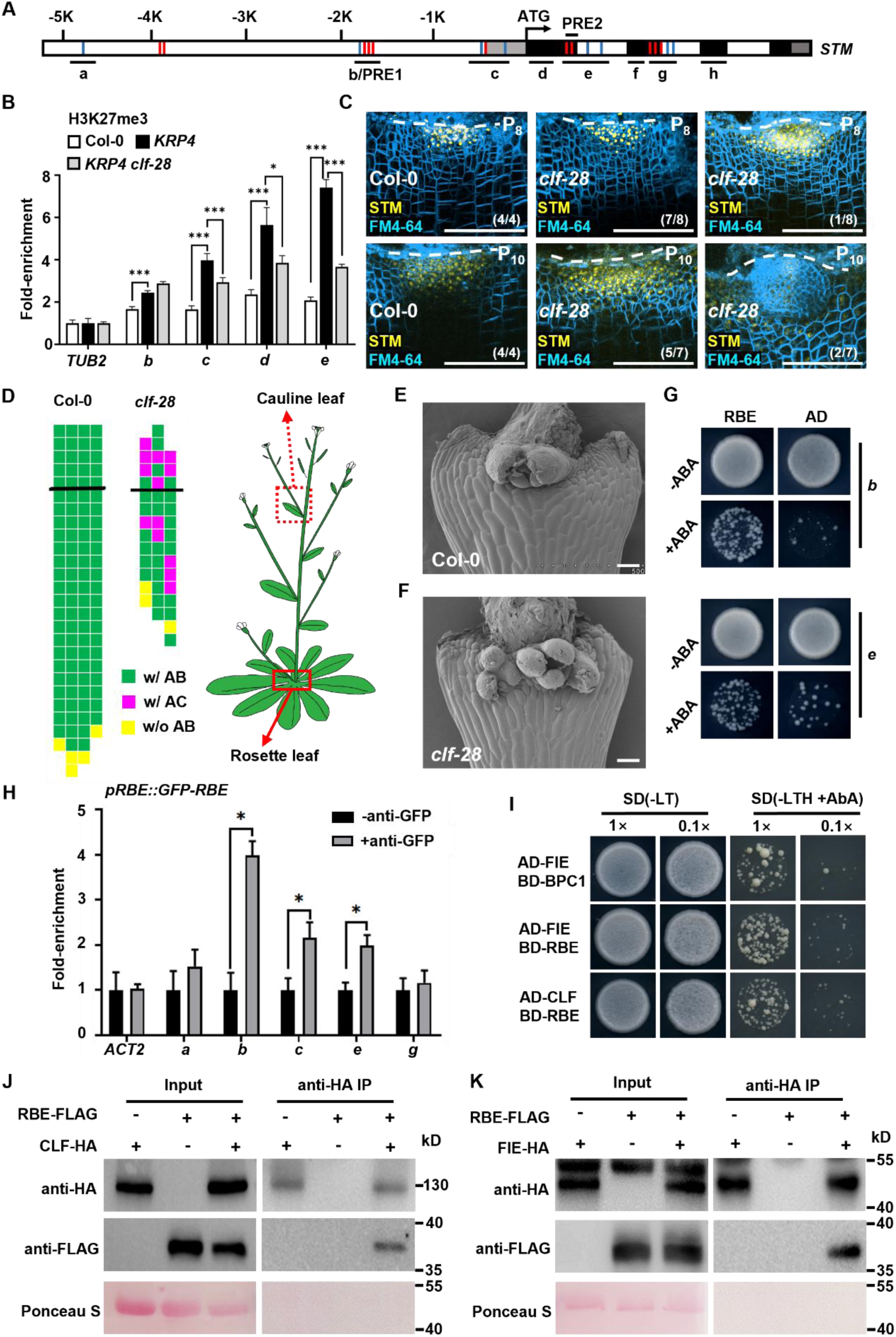
Silencing of *STM* expression in the leaf axils of *pCUC3:KRP4-mTQ2* is correlated with H3K27me3 and PRC2 complex. (**A**) Schematic diagram of the *STM* genomic region. ATG denotes the translation start site. The gray, black and white rectangles represent the 5’ untranslated region, coding region, and introns or intergenic regions, respectively. Each vertical red line indicates a GA repeat, and each vertical blue line indicates a telobox motif. The black solid lines represent the DNA fragments used in the ChIP and yeast one-hybrid assays. (**B**) Comparison of H3K27me3 levels at the *STM* locus in P_12-15_ leaf axil tissues of Col-0, *KRP4*, and *KRP4 clf-28*. The results from three representative biological replicate are shown, with three biological replicates showing consistent trends. The error bars represent the SDs of three biological replicates, and each run was performed in triplicate. (**C**) *STM* expression in the P_8/10_ leaf axils of Col-0 and *clf-28*. Reconstructed view of the leaf axil from short-day-grown plants showing the STM signal (yellow) and FM4-64 staining (cyan). The white dashed lines mark the incision line. Notably, a portion of *clf-28* plants (right column) presented substantially increased STM expression and faster AM initiation. (m/n) indicates that m in n biological repeats shows the displayed features. Scale bars, 100 µm. (**D**) Schematic diagrams of axillary bud formation in Col-0 and *clf-28*. Each column represents an individual plant, and each square represents a leaf axil. The thick black horizontal line denotes the border between the youngest rosette leaf and the oldest cauline leaf. The bottom row represents the oldest rosette leaf axils, with progressively younger leaves above. Green indicates the presence of a single axillary bud (w/ AB), yellow indicates the absence of an axillary bud (w/o AB), and magenta indicates the presence of one or more extra accessory bud(s) (w/ AC). Note the schematic of the shoot system of *Arabidopsis* showing the positions of rosette and cauline leaves (Right image). (**E** and **F**) Axillary buds in detached P_12-15_ leaf axils of Col-0 (E) and *clf-28* (F) after 5 d under short-day growth conditions. Plants were grown under short-day conditions for 14 days before in *vitro* culture. Note the formation of two buds in *clf-28*. Scale bars, 100 µm. (**G**) Y1H assay showing the interaction of RBE with the *STM* locus fragments, as shown in Fig. 2A. The transcription factors RBE were fused to the GAL4 activation domain (AD). AD alone served as a negative control. (**H**) ChIP assays verifying the interaction of RBE with *STM* locus fragments, as shown in Fig. 2A. An anti-GFP antibody was utilized, with ‘‘-anti-GFP’’ serving as a negative control. The error bars indicate the SDs of three biological replicates, and each run was performed in triplicate. *p < 0.05 (Student’s *t* test). (**I**) Y2H assay verifying the interaction of FIE and CLF with RBE and BPC1, with RBE and BPC1 used as bait proteins fused to the DNA binding domain (BD). And FIE and CLF fused to the GAL4 activation domain (AD). The left two columns represent the growth state of yeast on Leu/Trp-depleted media, and the two right columns represent the growth state of yeast on Leu/Trp/His-depleted and AbA-supplemented media. Sample dilution gradients of 1 × and 0.1 × were used. (**J** and **K**) Co-IP assays verifying the interactions between CLF and RBE (J), and between FIE and RBE (J). Total proteins were extracted from *Arabidopsis* leaf protoplasts coexpressing RBE-FLAG with CLF-HA, and RBE-FLAG with FIE-HA.

**Fig. 3.**
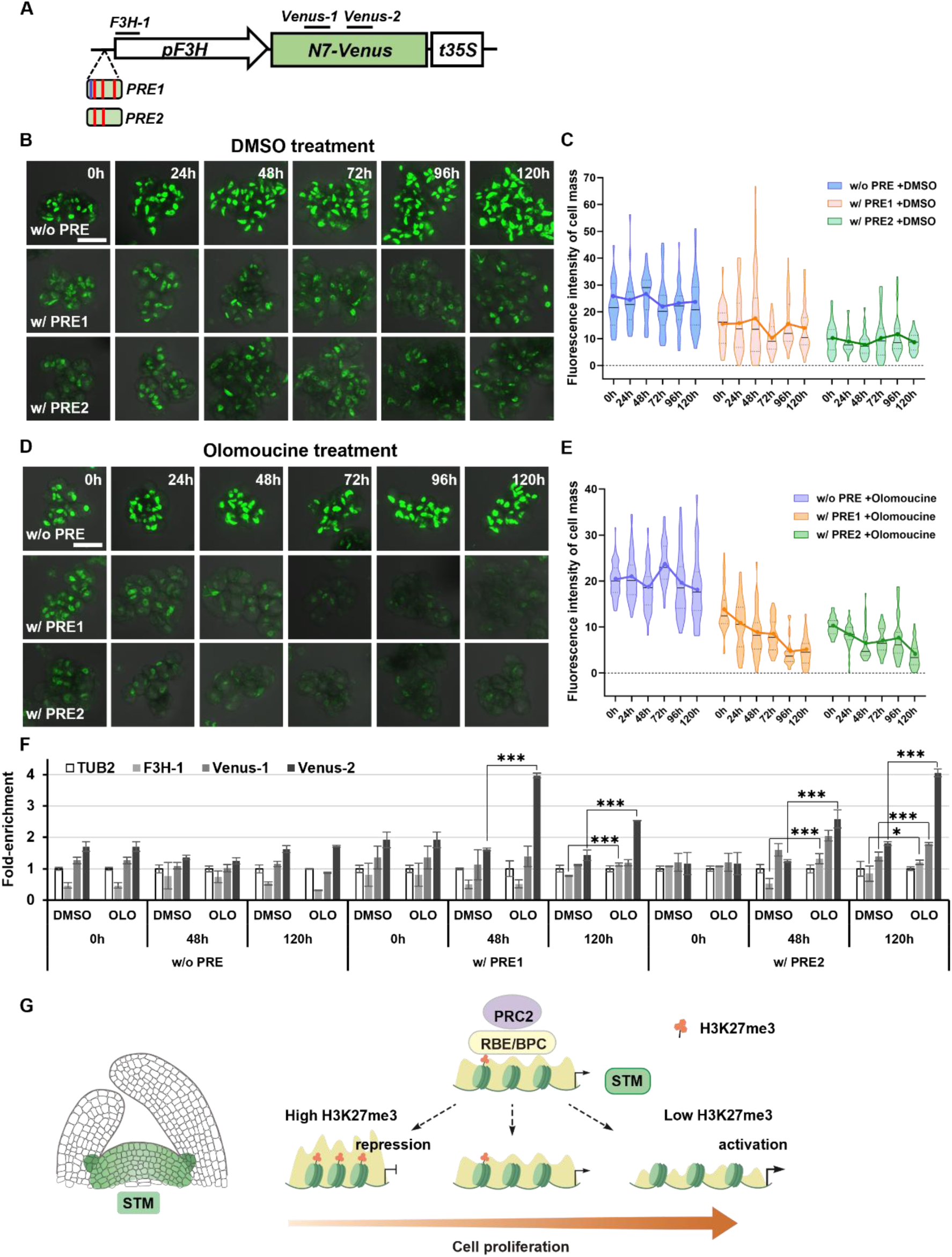
Cell division arrest results in H3K27me3 deposition. (**A**) Schematic diagram of the constructs used to test the relationships among cell division, H3K27me3 modification and gene expression. The promoter of the *F3H* gene serves as the basal constitutive promoter in this system. PRE1, containing telobox motifs (blue bars) and three GA repeats (red bars), was inserted upstream of the *F3H* promoter. PRE1 is from the base sequences of -2138 to -1931 before the *STM* gene start codon. PRE2, containing two GA repeats (red bars), was inserted upstream of the *F3H* promoter. The PRE2 element is located from the +214 to +399 base sequences after the start codon of the *STM* gene. *N7-Venus* is behind the *F3H* promoter. The cassette was integrated into T87 genome via the agrobacterium-mediated transformation. (**B** and **D**) *Venus* expression dynamics in different cell division states in *pF3H::Venus* without and after the addition of the PRE element. Cells with active (DMSO, B) and inactive divisions (olomoucine 100 µM, D) are shown. Scale bars, 50 µm. (**C** and **E**) Temporal changes in average fluorescence intensity of cell mass treated with DMSO (C) or olomoucine (E), with or without PRE elements. Observations and statistics were performed at chosen time points, including 0h, 24h, 48h, 72h, 96h and 120h. (**F**) ChIP-qPCR results showing a comparison of the effects of cell division on H3K27me3 accumulation. Three fragments, *F3H-1*, *Venus-1* and *Venus-2*, which are indicated by the short lines in (A), were tested. The error bars indicate the SDs of three biological replicates performed in triplicate. *p < 0.05, ***p < 0.001 (Student’s *t* test). (**G**) Conceptual model for the cell cycle-dependent maintenance of *STM* expression in AxP cells. PRC2 is constantly recruited to the *STM* locus to deposit H3K27me3. Cell proliferation antagonizes PRC2 to reduce H3K27me3 levels and drive *STM* expression.

In contrast, leaf axils of *pCUC3::CYCD3;1* plants exhibited enhanced *STM* expression levels (fig. S2, A and C). A high density of *STM*-expressing cells was observed in the P_9_ leaf axils of these plants (Fig. 1, P and Q), suggesting that accelerating the cell division rate promotes stem cell marker expression.

### Silencing of *STM* expression in leaf axils is correlated with increased H3K27me3

To investigate the mechanisms of *STM* silencing in non-dividing AxPs, we focused on epigenetic regulation due to its established role in cell fate transitions. Comparative analysis of histone modifications showed a significant increase in H3K27me3, which is linked to transcriptional repression, at the *STM* locus in *pCUC3::KRP4-mTQ2* leaf axil tissues compared to wild-type samples (Fig. 2, A and B).

We next examined whether H3K27me3 accumulation directly silences *STM* with the *clf-28* mutant, which lacks the PRC2 catalytic subunit *CURLY LEAF* (*CLF*) (*24*, *25*) required for H3K27 trimethylation (*26*, *27*). Live imaging showed that *clf-28* AxPs exhibited enhanced *STM* expression and accelerated AM initiation (Fig. 2C, and fig. S4, A to E). Additionally, *clf-28* plants developed extra accessory buds (Fig. 2D and fig. S4F), and a similar phenotype was observed in cultured detached leaves (Fig. 2, E and F and fig. S4G). Critically, introducing *clf-28* into *pCUC3::KRP4-mTQ2* rescued the axillary bud defects (fig. S4H) and reduced H3K27me3 levels in leaf axils (Fig. 2B). Together, these findings indicate that PRC2-mediated H3K27me3 deposition directly silences *STM*, thereby disrupting its maintenance in AxPs.

### RBE and BPCs recruit PRC2 to the *STM* locus

To further elucidate the mechanisms underlying H3K27me3 deposition, we investigated how PRC2 is recruited to specific genomic regions. Unlike in mammals, where PRC2 is directed to hypomethylated CpG-rich regions (*28*), plants employ sequence-specific recruitment via transcription factors including members from C2H2 zinc-finger (ZnF), APETALA2-like (AP2), and BASIC PENTACYSTEINE (BPC) families (*29*).

Our analysis of the *STM* locus identified putative Polycomb response elements (PREs), including 11 GA repeats and 8 teloboxes (A/GAACCCT/AA) (*29–32*), within its 5’ untranslated region (UTR) and coding region. These PREs are enriched in two regions, designated PRE1 and PRE2, with additional elements scattered elsewhere (Fig. 2A). The genomic region containing PRE1 is known to restrict *STM* expression (*33*). To investigate the function of PREs within the coding region, particularly those in PRE2, we generated a synonymous mutation, *mSTM*, by altering these PREs (fig. S5A). Expression of the *pSTM::mSTM-Venus* construct resulted in enhanced *STM* expression (fig. S5B) and development of enlarged lateral buds (fig. S5, D and E). Furthermore, in the *pCUC3::KRP4-mTQ2* background, this mutant version introduces weak but detectable *STM* expression, caused nuclei to maintain a rounded morphology instead of becoming fusiform (fig. S5C), and partially restored axillary bud formation (fig. S5D). Collectively, these findings underscore the role of PREs in repressing *STM* expression.

To identify putative PRC2-recruiting factors among boundary-expressed ZnF, AP2 and BPC members, we employed PRE-containing fragments as prey in a yeast one-hybrid screen. This screen identified RABBIT EARS (RBE), a ZnF protein, and BPC1 and BPC6, two homologous BPC proteins (Fig. 2G and fig. S4I), all exhibiting expression in the leaf axil (fig. S6, A to D), as potential modulators. Chromatin immunoprecipitation (ChIP) assays confirmed that RBE binds to three distinct telobox-containing regions within the *STM* locus (Fig. 2H). Although BPC1 was previously reported to bind the *STM* locus (*34*), we further mapped the specific binding sites of both BPC1 and BPC6 to the GA repeats within *STM* (Fig. 2A and fig. S4I).

Yeast two-hybrid (Y2H) assays demonstrated that RBE interacts with multiple PRC2 subunits, including FERTILIZATION INDEPENDENT ENDOSPERM (FIE), CLF, VERNALIZATION 2 (VRN2), SWINGER (SWN), and MULTICOPY SUPPRESSOR OF IRA1 (MSI1) (Fig. 2I and fig. S6E). These interactions between RBE and FIE or CLF were further validated by co-immunoprecipitation (co-IP) assays in leaf protoplast lysates (Fig. 2, J and K). Y2H also identified an interaction between BPC1 and FIE. In contrast, no direct physical interactions were detected between BPC6 and any PRC2 subunit. It is possible that BPC6 interacts directly with PRC1 (*32*), which in turn facilitates PRC2 recruitment.

We observed the formation of additional accessory buds in *rbe-2* loss-of-function mutant plants (*35*) (fig. S6, F and G), accompanied by upregulation of *STM* expression in the boundary region (fig. S6, H and I). Additionally, cell division rates were accelerated in the leaf axils of *rbe-2* (fig. S6J). Similarly, accessory bud formation and elevated *STM* expression were observed in *bpc1* and *bpc6* loss-of-function mutants, as well as in the *bpc1 bpc6* double mutant (fig. S6, F to H). These results demonstrate that RBE, BPC1 and BPC6 mediate *STM* silencing in AxPs through the recruitment of PRC2.

### Division eliminates H3K27me3 and maintains expression

The connection between *STM* expression and cell proliferation suggests a mechanism involving replication-coupled dilution of H3K27me3 at the *STM* locus. To test this hypothesis, we adopted an *Arabidopsis* cell culture system featuring olomoucine-induced division arrest (fig. S7) (*36*). Transcriptome data indicated that PRC2 components, along with the recruiting factors RBE, BPC1 and BPC6, are expressed in the cells (*37*). We introduced a *Venus* reporter driven by the constitutive *pF3H* promoter, which is free of H3K27me3 (Fig. 3A), leading to detectable fluorescence in both proliferating and arrested cells. However, when PRE regions bound by RBE and BPC were fused to the *pF3H* promoter, we observed a reduction in *Venus* signal and the deposition of H3K27me3 (Fig. 3, B, D and F). H3K27me3 levels also notably increased following division arrest (Fig. 3F). Consistent with this proliferation-associated increase in H3K27me3, live imaging showed that Venus signals gradually declined in arrested cells (Fig. 3, D and E) but were maintained in proliferating cells (Fig. 3, B and C, and fig. S8). Collectively, these findings indicate that cell division dilutes PRE-recruited H3K27me3, thereby sustaining gene expression (Fig. 3G).

In parallel, we utilized a mesophyll protoplast culture system to independently validate the influence of cell proliferation on gene expression maintenance (fig. S9). This approach took advantage of the natural heterogeneity in proliferation among cultured mesophyll cells (*38*, *39*). By targeting *pF3H::Venus* to a single genomic locus via Cre/loxP recombination, we minimized positional effects. The incorporation of each PRE resulted in elevated H3K27me3 levels (fig. S9E). In agreement with the cell culture results, Venus signals diminished in non-dividing cells but remained stable in actively dividing cells when the PRE was present (fig. S9, A to D), further supporting a cell cycle-dependent mechanism of epigenetic dilution.

### A cell cycle-dependent epigenetic Sisyphus model for stem cell maintenance

Our findings reveal that both PRE-dependent H3K27me3 deposition and cell proliferation modulate epigenetic states to balance *STM* expression and AxP differentiation. To formalize these regulations, we developed a hybrid mathematical model integrating histone modification dynamics (*40–43*), *STM* expression, and cell division (Fig. 4A, fig. S10A, Table S1, and Methods). The model is built upon three core principles:

1. PRC2, recruited by RBE and BPC1, continuously deposits H3K27me3 at the *STM* locus (Fig. 2, G to K and fig. S4I). This process includes autocatalytic propagation of H3K27me3 (*25*), along with stochastic methylation and demethylation events.
2. *STM* transcription is inversely correlated with H3K27me3 levels. *STM* transcription actively contributes to the removal of this mark (*44*, *45*), and STM engages in self-activation (*8*), while protein degradation helps maintain homeostasis.
3. Cell division dilutes H3K27me3 (Fig. 3, A to E and fig. S9, A to E) through nucleosome segregation (*46*), with slower proliferation rates accelerating epigenetic silencing (Fig. 2, A and B).

**Fig. 4.**
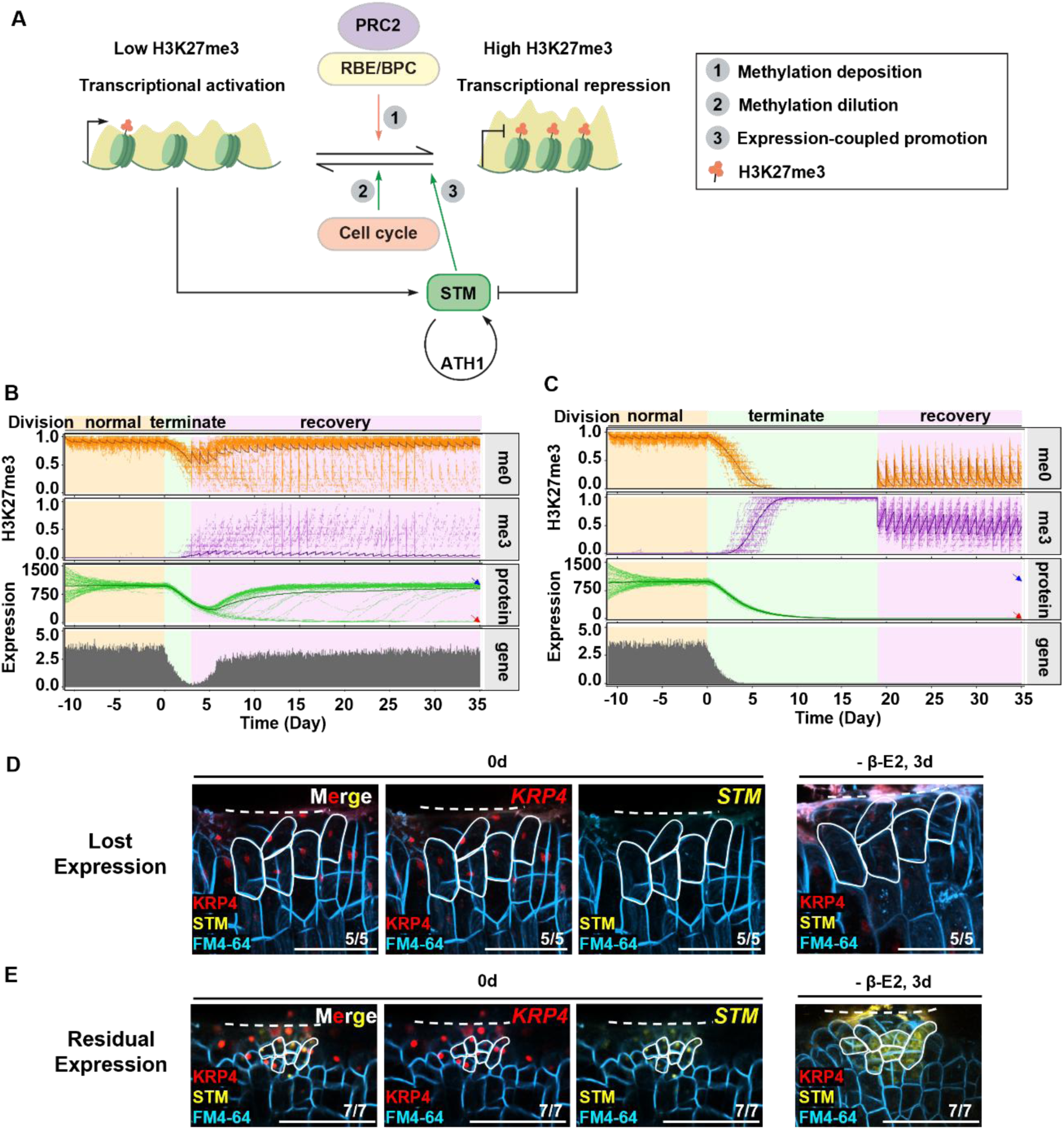
The hybrid histone modification-*STM* expression-cell division model and its prediction and validation. (**A**) Schematic of the model showing key feedback mechanisms. Active chromatin has little or almost no H3K27me3 (left state), whereas repressed chromatin has more H3K27me3 deposition (right state). The repressive state, characterized by H3K27me3 (orange pentagon), positively recruits PRC2 via the transcription factors RBE and BPCs (orange arrow). The active chromatin state can be achieved through cell cycle progression and *STM* expression (green arrows), and *STM* self-activation mediated by the ATH1‒STM complex can maintain continuous *STM* expression (black circular arrow). The numbers 1, 2 and 3 represent three molecular processes—methylation deposition, methylation dilution, and expression-coupled promotion—with the latter two processes functioning together to reduce H3K27me3 levels. The black two-way unilateral arrows represent state transitions; the colored arrows represent feedback interactions. The black arrows represent transcriptional activation, and the black symbols represent transcriptional repression. For clarity, not all model elements are shown. See fig. S6A for a more elaborate schematic. (**B** and **C**) Representative trajectories from stochastic simulations of 3-d (B) and 19-d (C) cell division arrest before recovery. Simulations were started randomly from ±50% fluctuation of protein stable points in pluripotent cells (uniform H3K27me0) with a normal cell cycle (22 h), and 12 cell cycles were performed for preequilibration before quiescence induction. Each graph shows 80 overplotted trajectories. The thick line in each panel represents the average curve of the cell population. Red arrowhead, differentiated group; blue arrowhead, pluripotency-restored group. Gene activity was measured as the average number of transcription events in the cell population per 15 min interval. The other parameters are fixed to their default values, as shown in Table S1. (**D‒E**) *STM* expression and cell division in P_13/14_ leaf axils of *pCUC3::XVE pLexA::KRP4-mTQ2* plants grown under short-day conditions for 18 d with 10 µM *β*-estradiol (*β*-E2). Leaves were detached (left, 0 d) and cultured for 3 d without *β*-E2 (right, - *β*-E2, 3 d). The expression of *STM* (yellow) and *KRP4* (red) is shown. (D) Leaf axil with fully lost STM signals. (E) Leaf axil with remnant STM signals at 0 d. Note that the STM signal was undetectable in (D) by 3 d. Selected cells were white-delineated for easy tracking of *STM* expression and cell division. (m/n) indicates that m in n biological repeats shows the displayed features. The white dashed lines mark the incision line. Scale bars, 50 µm.

The model generates sigmoidal protein production dynamics coupled with linear degradation, enabling two distinct stable states: a pluripotent state characterized by low H3K27me3 and active *STM* expression, and a differentiated state with high H3K27me3 and silenced *STM* (fig. S10B). Parameter analysis confirmed robust bistability at physiological division rates (fig. S10, D, F and G), whereas altered cycling frequencies disrupted bistability (fig. S10, C and E), indicating a unimodal dependence on the cell cycle. Computational simulations aligned closely with experimental observations: accelerated division upregulated *STM* expression (fig. S10H), while decelerated division led to progressive *STM* downregulation and promoted differentiation in subpopulations of cells (fig. S10I, Methods).

After establishing concordance between the model and experimental data (Fig. 4A and fig. S10A), we assessed additional model parameters. Introducing variation in cell-cycle arrest duration revealed that division suspension reduced *STM* expression, with prolonged quiescence increasing the probability of irreversible differentiation (Fig. 4, B and C and fig. S11, A to H), resulting in a greater number of irreversibly differentiated cells.

We further modeled potential strategies for rescuing pluripotency. Simulated disruption of H3K27me3 successfully restored pluripotency (Fig. 5A), an effect attributable to stochastic transcription and *STM*-mediated positive feedback. In contrast, simply enhancing *STM* self-activation—for example, via *ATH1* overexpression (*8*)—failed to restore stem cell identity due to the stability of locus silencing (Fig. 5B, Methods).

**Fig. 5.**
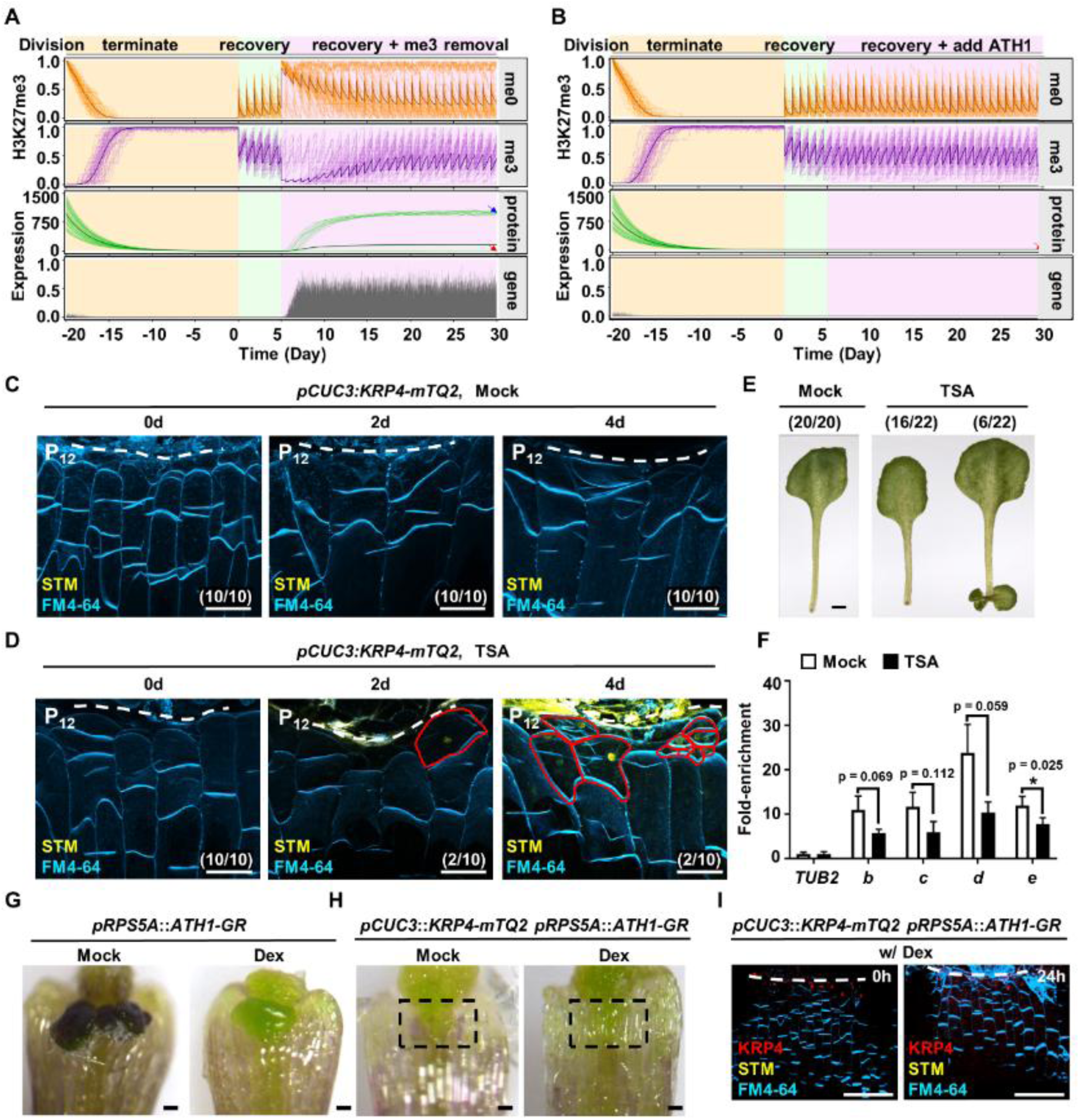
TSA treatment but not ectopic expression of ATH1 enables re-expression of *STM*. (**A** and **B**) Representative trajectories from stochastic simulations of (A) H3K27me3 removal and (B) *ATH1* overexpression after long-term quiescence. The simulations were started randomly from ±50% fluctuation of protein stable points in quiescent pluripotent cells (uniform H3K27me0), after which division was stopped for 20 d, and the cells were then allowed to rest for another 5 d before treatment. In (A), each histone in the simulation samples were reset to H3K27me0 with a probability of 95%. In (B), the maximum self-activation level of the *STM*, 𝜀, was increased by 20%. Each graph shows 80 overplotted trajectories. The thick line in each panel represents the average curve of the cell population. Red arrowhead, differentiated group; blue arrowhead, pluripotency-restored group. Gene activity was measured as the average number of transcription events in the cell population per 15 min interval. The other parameters are held at their default values, as shown in Table S1. (**C** and **D**) Time-lapse imaging of detached P_12_ leaf axil cells of *pCUC3::KRP4-mTQ2* with mock (C) or TSA (D) treatment. *STM* (yellow) expression was reactivated in a portion of leaf axil cells (red solid line boxed) after TSA treatment but not after mock treatment. (m/n) indicates that m in n biological repeats shows the displayed features. Scale bars, 50 µm. (**E**) Axillary bud initiation in *pCUC3::KRP4-mTQ2* leaf axils after TSA but not mock treatment. (m/n) indicates that m in n biological repeats shows the displayed features. Scale bars, 1 mm. (**F**) Comparison of H3K27me3 levels at the *STM* locus in the leaf axils of P_12-16_ in *pCUC3::KRP4-mTQ2* after mock or TSA treatment. The error bars indicate the SDs of three biological replicates, and each run was performed in triplicate. *p < 0.05 (Student’s *t* test). (**G** and **H**) Bud formation after mock or Dex treatment in *pRPS5A::ATH1-GR* (G) and *pCUC3::KRP4-mTQ2 pRPS5A::ATH1-GR* (H) plants. Note the enlarged meristem in (G), and lack of axillary bud formation (black dashed boxed region) even after Dex treatment in (H). Scale bars, 100 µm. (**I**) Expression of *STM* in the *pCUC3::KRP4-mTQ2 pRPS5A::ATH1-GR* leaf axil before and after Dex treatment. Note that Dex treatment was not able to restore STM signals (yellow). KRP4 is shown in red, and the plasma membrane was stained with FM4-64 (cyan). Scale bars, 50 µm.

### Prolonged quiescence switches AxPs to irreversible differentiation

Mathematical modeling leads to testable predictions. We first validated the bistable *STM* expression dynamics, in which prolonged quiescence triggers irreversible silencing (Fig. 4, B and C and fig. S11, A to H). We introduced an inducible *pCUC3::XVE pLexA::KRP4-mTQ2* into *pSTM::STM-Venus* plants. Induction of *KRP4-mTQ2* gradually diminished STM-Venus signals. Importantly, cells that had completely lost STM-Venus signal failed to restore it even after the termination of *KRP4-mTQ2* expression (Fig. 4D). In contrast, cells retaining residual STM-Venus signal resumed division following *KRP4-mTQ2* termination (Fig. 4E). These results experimentally confirm that the bistable switch prevents *de novo* activation of pluripotency.

Consistent with model predictions that epigenetic perturbation could reactivate *STM* (Fig. 5A), treatment with trichostatin A (TSA)—a chemical inhibitor that promotes H3K27 acetylation and inhibits methylation (*47*)—reduced H3K27me3 levels at the *STM* locus (Fig. 5F), restored *STM-Venus* expression in a subset of leaf axil cells (Fig. 5, C and D), and induced sporadic axillary buds in *pCUC3::KRP4-mTQ2* plants (Fig. 5E). Conversely, enhancing *STM* activation via dexamethasone-induced *pRPS5A::ATH1-GR* failed to restore axillary buds (Fig. 5, B and G to H) or STM-Venus signals in quiescent AxPs (Fig. 5I). These results further corroborate the model’s predictions.

## Discussion

Cell fate is intrinsically linked to cell cycle dynamics, especially in stem cells. While mammalian studies have extensively explored how proliferation influences pluripotency (*2*)—with adult stem cells often maintaining their fate through quiescence (*4*, *48–51*)—the mechanisms in plants remain less understood. Here, using AxP cells (*7*) as a model, we demonstrate that slowed proliferation paradoxically accelerates differentiation, contrasting with mammalian paradigms. Conversely, accelerating proliferation enhanced meristem formation and reinforced pluripotency (Fig. 1, fig. S1 and fig. S2). This aligns with previous reports that repression of boundary genes enables cell cycle activation and promotes AM initiation (*52*).

The bistable regulation of *STM* depends on opposing epigenetic forces: sequence-specific H3K27me3 deposition silences *STM* and drives differentiation, while proliferation dilutes this mark to sustain expression. Although DNA replication-coupled H3K27me3 maintains mechanisms exist (*53*), maintenance is not always complete (*46*). Incorporating *STM*’s self-activation loop (*8*), we propose a cell cycle-coupled “Sisyphus” mechanism in which targeted H3K27me3 deposition and replication-dependent dilution govern fate bistability (Fig. 4A and fig. S10A). Computational and experimental evidence confirms that prolonged quiescence irreversibly silences *STM* through stable epigenetic modification, locking cells into differentiation (Fig. 4, C and D).

Beyond pluripotency maintenance, proliferation-mediated dilution of H3K27me3 may broadly regulate cell fate transitions. For example, floral termination involves similar dilution at differentiation-promoting loci like *KNUCKLES* (*36*, *54*). Plants uniquely employ sequence-dependent PRC2 recruitment via PRE-bound transcription factors (*29*, *31*), enabling precise locus-specific tuning of H3K27me3. In contrast, mammals—which lack consensus PREs—rely on sequence-independent PRC2 recruitment (*28*) and bivalent chromatin domains (e.g., H3K27me3/H3K36me coexistence) (*55*) for fate plasticity. These differences highlight an evolutionary divergence that underscores proliferation-coupled epigenetic regulation as a plant-specific innovation.

## Acknowledgments

We are grateful to Drs. Chuck Gasser, Arp Schnittger, Hong Wang, Tengbo Huang, and Ping Yin for plant materials and expression vector. We thank the Core Facilities of Life Sciences, Peking University, and the State Key Laboratory of Plant Genomics for assistance with SEM.

## Author contributions

Conceptualization: YJ, YW

Design: YJ, YW, RS

Molecular genetics, phenotyping, and image analysis: YG, BS, HW

Computational modeling: YY

Computational modeling supervision: YJ, LT

Contributed experimental reagents: YH, MU

Contributed PRC2 interacting TF data: JX, DW

Writing – original draft: YJ, YW, YG, YY

Chart Layout and Chart Design: YJ, YW, YG, YY

## Materials and Methods

### Experimental model and subject details

*Arabidopsis thaliana* plants were grown in soil under constant light at 22°C. For AM initiation genotyping, plants were grown under short-day conditions (8 h light and 16 h dark at 22°C) for 30 d and then under long-day conditions (16 h light and 8 h dark at 22°C) for 30 d to induce flowering before the axillary buds were counted. For live imaging of leaf axil regions, plants were grown under short-day conditions for 15 d or 21 d. For gene expression profile analysis, plants were grown under short-day conditions for 15 d or 21 d before being transferred to low-melting-point agarose for sectioning. For mRNA quantitative analysis and ChIP assays, plants were grown under long-day conditions for 14 d (with seedlings).

### Genetic material

The *Arabidopsis thaliana* ecotypes Columbia (Col-0) and Landsberg *erecta* (L*er*) were used as wild-type plants. The *krp3467*, *krp6*, *krp47*, *krp3*, *krp5*, *krp7* (*56*), *ath1-3* (SALK_113353) (*57*), *clf-28* (SALK_139371) (*58*), *bpc1* (SALK_072966) (*59*), *bpc6* (SALK_055387) (*59*), *pSTM::STM-Venus* (*60*), *pWUS::3×Venus-N7* (*61*), *rbe-2* (SALK_037010) (*62*), *pRBE::RBE-GFP* (*62*), and *pRPS5A::ATH1-GR* (*8*) strains were established in the Col-0 background. *p35S::STM-GR* (*63*) is in the L*er* background. The genotyping primers used are listed in Table S2.

## Method details

### Construction of transgenic plants

For *pCUC3::KRP4-mTurquoise2* (*pCUC3::KRP4-mTQ2*) construction, we used the GreenGate system to assemble the *pCUC3* promoter (4358 bp upstream of the start codon), full-length *KRP4* CDS, mTQ2 sequence, and terminator (pGGE001) into the destination vector pMOA34 to confer hygromycin resistance.

For *pCUC3::CYCD3;1* construction, we used the GreenGate system to assemble the *pCUC3* promoter (4358 bp upstream of the start codon), full-length *CYCD3;1* CDS, and terminator (pGGE001) into the destination vector pMOA34 to confer hygromycin resistance.

For *the pCUC3::XVE pLexA::KRP4-mTQ2* construct, we also used the GreenGate system to assemble the *pCUC3::XVE and pLexA::KRP4-mTQ2 components and then transferred these two components together into the destination vector pMOA34* to confer hygromycin resistance.

For *the pSTM::mSTM-Venus* construct, we used the homologous recombination system to assemble the *pSTM* promoter (5736 bp upstream of the start codon), full-length mutanted-*STM*-CDS, Venus sequence and OCS terminator into the destination vector pMOA36 to confer Basta resistance.

### Chemical treatment and RT-qPCR

For Dex treatment, a 10 mM stock solution of Dex (Sigma-Aldrich) in ethanol was diluted with distilled water or 1/2 MS medium to a final concentration of 10 μM for treatment of the leaf axils. Water with only ethanol was used as a mock control. For β-estradiol treatment, a 20 mM stock solution of β-estradiol (Sigma-Aldrich) in DMSO was diluted with distilled water to a final concentration of 10 μM for treatment of the leaf axils. For gene expression analyses, plants were grown for 14 d under short-day conditions, and meristematic and boundary-enriched tissues were manually dissected by removing the leaves and roots. Total RNA was extracted with the AxyPrep Multisource RNA Miniprep Kit (Corning). First-strand cDNA synthesis was performed via TransScript One-Step gDNA Removal and cDNA Synthesis SuperMix (TransGen). Quantitative reverse transcriptase PCR (RT-qPCR) was performed on a Bio-Rad CFX96 real-time detection system via the KAPA SYBR FAST qPCR Kit (KAPA Biosystems). The relative RT-qPCR expression was normalized to that of ACTIN2 (AT3g18780), which shows constant expression under various treatments and has been shown to be a superior reference gene for qRT-PCR analysis. Gene-specific primers (Table S2) were used to amplify and quantify each gene.

### Tissue preparation, confocal microscopy and scanning electron microscopy

Seedlings were grown in MS media under short-day conditions (8 h light and 16 h dark at 22°C) for 15 d or 21 d after seed germination. Leaves between P_8‒16_ were detached from the seedlings for live-imaging analysis. For low-melting-point agarose sections, the shoot apices of 15-d-old or 21-d-old short-day-grown seedlings were collected. The samples were either live-imaged or fixed and sectioned as previously described (*64*). Optical photographs were taken with a Nikon SMZ1000 stereoscopic microscope or an Olympus BX60 microscope equipped with a Nikon DS-Ri1 camera. Confocal microscopy images were taken with a Nikon A1 confocal microscope. The excitation and detection wavelengths for GFP, Venus, CFP, mTQ2, TagRFP, propidium iodide (Sigma-Aldrich), FM4-64 (Thermo Fisher), and autofluorescence were as previously described (*9*, *64*). The detection conditions for the Fluorescent Brightener 28 (FB28, Sigma-Aldrich) staining reagent were identical to those for CFP. For the axillary bud phenotype after 3‒5 d *in vitro*, scanning electron microscopy was performed using a Hitachi S-3000N variable pressure scanning electron microscope.

### EdU incorporation assay

Seedlings were germinated and then grown in MS media under short-day conditions for 15 d. Meristematic and boundary-enriched tissues (with the leaves and hypocotyls removed) were placed in 1/2 MS liquid media supplemented with 10 μM EdU for 6 h and then fixed in 3.7% formaldehyde for 15 min at room temperature. The samples were then washed twice with 3% bovine serum albumin and permeabilized with 1% Triton X-100 for 20 min. The EdU-labeled nuclei were labeled with an Invitrogen Click-iT® EdU Imaging Kit (C10337). Confocal microscopy images were taken using a Nikon A1 confocal microscope. To detect the Alexa Fluor 555 signal, a 561-nm laser was used for excitation, and the emission was measured at 570 to 620 nm.

### Time-lapse imaging

Time-lapse imaging of leaf axils was performed using a Nikon A1+ confocal laser scanning microscope equipped with a 40× or 60× water-immersion objective. Leaves between P_8_ and P_10_ were detached from the seedlings for live-imaging analysis. If necessary, 10 μg/mL FM 4-64 or 100 μg/mL PI was applied to the dissected leaf axil for 10 min. The leaf axil was then mounted in imaging medium (1/2 MS medium topped with 1% agarose, w/v) and submerged in water for imaging. For time-lapse experiments, sterile water was used for imaging, and the dissected leaf axil was transferred to a new growth medium and cultured *in vitro* for 24 h under normal growth conditions after each imaging session.

### Image analysis

Confocal images were converted to TIFF format using FIJI (*65*) and then processed using MorphoGraphX (MGX) following the User Manual (*66*). The mesh was smoothed and subdivided into at least 200,000 vertices. The cell contours were projected onto the extracted surface at a depth of 1‒3 μm from the epidermis.

For analysis of cell growth, as shown in Fig. 1 and fig. S1, the cell contours were stained with FM4-64. The cells were segmented manually, and the cell lineage for the corresponding time interval was determined by matching the mother and daughter cells. Cell identity was determined on the basis of the adaxial-abaxial markers expressed at the previous time point because these markers may change during growth of the primordia. The cell lineage (Fig. 1, E, G, and I) results were used to determine the cellular growth rate, and number of cell divisions. The cell growth rate (fig. S1, A to C) was defined as the ratio of the cell areas in consecutive measurements, i.e., the current cell area divided by the cell area at the previous time point. The number of cell divisions was counted using the cell lineage maps. Heatmaps of the cell growth rate for each interval are displayed at the first of the two time points.

### Chromatin immunoprecipitation (ChIP)

ChIP experiments were performed according to previously published protocols (*67*). The meristematic and boundary-enriched tissues of 14-d-old short-day-grown *pRBE::GFP-RBE* plants were collected for ChIP with an anti-GFP antibody (11814460001, Roche). No-antibody controls were used to calculate enrichment. PCR was performed using the precipitated DNA as a template. DNA fragment enrichment was determined by quantitative PCR analysis. Three independent biological replicates were performed for each ChIP analysis.

Ultra-low-input native ChIP (ULI-NChIP) and qPCR analyses were performed according to published protocols with modifications (*10*, *68*). More than 100 leaf axil tissues (basal 1 to 3 mm) were dissected from late immature stage (P_12_ to P_16_) leaves for each replicate. The tissues were fully ground in 30 μL of Galbraith buffer (45 mM MgCl_2_, 30 mM sodium citrate, 20 mM MES, 0.5% Triton X-100, pH 7.0). The pestle was washed with an additional 30 μL of Galbraith buffer into the same tube. Nuclei were centrifuged at 1000 × *g* for 10 min at 4°C. The supernatant was discarded, and the sediment was resuspended in 50 μL of nuclear isolation buffer (NUC-101; Sigma-Aldrich). The chromatin was digested with 100 units of micrococcal nuclease (M0247; New England Biolabs) at 37°C for 7.5 min, 10% of the sample was removed as input, and then the sample was precleared with 10 μL of 1:1 protein A:protein G Dynabeads (Life Technologies). Chromatin was immunoprecipitated with 1 μg of anti-H3K27me3 (07-449; Merck) in antibody-bead complexes at 4°C overnight. Protein‒DNA‒bead complexes were washed twice with 400 μL of low-salt wash buffer (20 mM Tris-HCl; pH 8.0; 2 mM EDTA;150 mM NaCl; 1% Triton X-100; and 0.1% SDS) and twice with high-salt wash buffer (20 mM Tris-HCl; pH 8.0; 2 mM EDTA; 500 mM NaCl; 1% Triton X-100; and 0.1% SDS). The protein‒DNA complexes were eluted in 30 μL of newly prepared ChIP elution buffer (100 mM NaHCO_3_ and 1% SDS) at 65°C for 90 min. Fragmented DNA was purified using phenol/chloroform/isoamyl alcohol, precipitated with ethanol and finally dissolved in 20 μL of elution buffer. All primers used for ChIP-qPCR are listed in Table S2.

### Quantification and statistical analysis

For AM initiation analysis (as shown in Fig. 2D, fig. S1E, S1F, S4H, S5D and S6F), we performed at least two independent biological experiments, and more than three individual plants were counted each time. For the detached leaf cultures (as shown in Fig. 1, K to M, and Fig. 2, E and F, and fig. S6G), we performed two independent biological experiments in which more than ten individual leaves were cultured for each stage.

For RT-qPCR and ChIP-qPCR analysis (as shown in Fig. 2B, 2H, 3F, 5F and fig. S4E, S6H and S9E), at least three independent biological experiments were performed, and each experiment was run in triplicate. Statistical analysis was performed through Student’s *t* test in Excel, and P values less than 0.05 were considered to indicate significant differences for any set of data. For all qPCR data, the 2^-ΔΔCT^ method was applied following sequential normalization to input and housekeeping genes.

### Yeast one-hybrid assay

The yeast one-hybrid assay used the Y1H Gold strain. The *STM* bait sequences were inserted into *pAbAi*. The full-length coding sequences of BPC1, BPC6 and RBE were fused with the GAL4 DNA binding domain in *pMDEST22*. Yeast cells were grown on media lacking Ura, Trp, and adenine (A) to test for protein‒DNA interactions. Yeast cells were plated on nonselective synthetic defined media lacking Ura and Trp to successfully transform the plasmids.

### Yeast two-hybrid assay

The yeast two-hybrid assay used the Y2H Gold strain. The full-length coding sequences of CLF, FIE, VRN2, SWN, and MSI1 were fused with the GAL4 activation domain in *pMDEST32*. The full-length coding sequences of BPC1 and RBE were fused with the GAL4 DNA binding domain in *pMDEST22*. Yeast cells expressing the GAL4 DNA-binding domain and activation domain-fused proteins were grown on stringent selective synthetic defined (SD) medium lacking Trp, Leu, His, and adenine to test for protein interactions. Yeast cells were plated on nonselective SD medium lacking W and L to identify successful transformants.

### Coimmunoprecipitation (co-IP)

To express tag-fused proteins in *Arabidopsis* leaf protoplasts, the full-length coding sequences of CLF and FIE were fused with HA in *pUC19-p35S::HA*; the full-length coding sequence of RBE was fused with FLAG in *pUC19-p35S::FLAG*. The constructs were coinfiltrated into *Arabidopsis* leaf protoplasts. Immunoprecipitation experiments were performed after overnight incubation, and controls were used to express only the FIE or CLF proteins or only the RBE protein. The samples were collected, and proteins were extracted with extraction buffer (50 mM HEPES [pH 7.5], 150 mM NaCl, 1 mM EDTA [pH 8.0], 0.5% Triton X-100, 20 mM 50× cocktail inhibitor (Roche), 10 mM DTT, and 1 mM PMSF). After centrifugation, the supernatant was incubated with anti-HA beads (MBL, 561-8). Tagged proteins were detected with anti-FLAG (Sigma-Aldrich, Cat# F1804) and anti-HA (CST, C29F4) antibodies.

### Agrobacterium-mediated transformation of *Arabidopsis* T87 cells

The *Arabidopsis* T87 suspension cell line and the *pPLink* binary expression vector, gifts from Prof. Ping Yin, were used for gene expression in this study. Transformation, selection, and culture of the transgenic T87 suspension lines were performed according to previously published protocols (*36*, *69*). Transgenic T87 calli, post-screening and verification, were subjected to suspension culture for two weeks. Subsequently, the samples were harvested for ChIP assay and confocal microscopic imaging. For the ChIP assay, approximately 200 µL of pelleted cells (dry weight) were collected by gravitational sedimentation. For expression assay of the reporter lines, confocal microscopy images were taken with a Nikon A1.

The Venus fluorescence intensity within cell mass was quantified using ImageJ. For each line and time point, a minimum of 30 independent cell mass were analyzed, as shown in Fig. 3, C and E. The corrected Mean Fluorescent Intensity of Cell mass (MFIC), representing the actual Venus fluorescence intensity per cell mass, was calculated using the following formula: MFIC= [Integrated Density - (Area of Venus signal × Mean fluorescence of background)] / Area of Venus signal.

### Computational model and simulation

We constructed a model of the biological process at the *STM* locus to explain the experimental results by expanding upon a previous mathematical model of a generic PRC2 target gene (*40*, *43*). The model considers chromatin histone modification, the *STM* expression cycle, and DNA replication (fig. S5D). An exhaustive description of the methods used, together with the code, can be found at https://github.com/yy420106/stmExpr-chromMod-cellDivModel. The optimal model parameter values for determining the observed primordia morphologies are listed in Table S1.

## Supplementary Text

### Computational Methods and Simulation Details

#### Mathematical Model

##### Overview

We constructed a biological process model of the *SHOOT MERISTEMLESS* (*STM*) gene locus to explain our experimental results by expanding a previous mathematical model of a generic polycomb target gene (*40*, *43*). The new model is composed of three main inter-coupled sections, i.e., chromatin histone modification, *STM* expression cycle and DNA replication/cell division (Fig. 4A and fig. S10A). The dynamics of H3K27 methylation, demethylation, gene transcription and translation, protein degradation, as well as DNA replication were simulated using the Gillespie’s stochastic simulation algorithm (SSA) (*70*), a classical and widely used method for simulating the stochastic behavior of chemical systems against time evolution by Monte Carlo sampling. This algorithm is defined by a series of possible reactions or events, and a corresponding propensity for each event to occur (master equation). At each iteration, the time interval Δ𝑡 for the next coming event and the event itself are both selected probabilistically based on Poisson process. Subsequently, the system is updated by performing the selected event, and time is incremented by Δ𝑡.

In our simulations, we tracked the methylation status at H3K27, denoted by 𝐻_𝑖_, of each H3 histone 𝑖 ∈ [1, 2 … , 𝑁_h_] within the *STM* locus on chromatin, where 𝐻_𝑖_ ∈ {me0, me1, me2, me3} represents the number of methyl group. Here we set 𝑁_h_ = 36 to approximate the physical length of *STM* gene (3482 bp) and its promoter, according to the putative value that a nucleosome contains ∼200 bp (include linkers). Histones with the indices 2𝑗 − 1 and 2𝑗, for each positive integer 𝑗, were thought to be in the same nucleosome. Meanwhile, we recorded the number of STM protein molecules (actually we did not restrict its values to integers, discuss later), 𝑁_p_ in each time step to characterize the *STM* expression level and stem cell fate. Overall, we considered totally 2𝑁_h_ + 2 possible events ( 𝑁_h_ histone methylations, 𝑁_h_ histone demethylations, gene expression and protein degradation) in our model. Note that, not all events are possible at any time, e.g., methylation of trimethylated histones or degradation when proteins are no longer presented. The event propensities, which are elaborated in the following text, as denoted by 𝑟(with a specific superscript), are recalculated after each system update. Additional information about model parameters is shown in Table S1.

##### Chromatin Histone Modification

***H3K27 methylation.*** In our model, polycomb repressive complex 2 (PRC2) activity results in the methylation of H3K27 (Fig. 4A and fig. S10A). PRC2 is a large protein complex containing an enzymatic subunit with histone methyltransferase activity (*71–74*), and a non-catalytic subunit that recognizes H3K27me3/me2 (*75*). Therefore, PRC2 can recognize existing H3K27me3/me2 and then add methyl group to other histones via a read/write mechanism (*25*, *75*, *76*), forming a conserved mechanism in both animals and plants. Meanwhile, both *in vitro* and *in vivo* experimental evidence suggested that PRC2 mostly worked in a non-processive manner (*77*, *78*), adding one methyl group at a time, which we applied in our simulation.

To model H3K27 methylation, we followed the ideas of a previous work (*40*), but with slight differences. Overall, the propensity of methylation, 𝑟_i_^me^for histone 𝑖 , depends on the current methylation status of the surrounding histones, together with some unusual long-range interactions through DNA looping. Besides, 𝑟_i_^me^ also depends on two reference rates: the PRC2-dependent methylation rate 𝑘_me_ and stochastic methylation rate 𝛾_me_. Moreover, our model also takes the different catalytic abilities of PRC2 on specific methylated substrates (H3K27me0/me1/me2) into account, as supported by *in vitro* experiments (*77*). For 1 ≤ 𝑖 ≤ 𝑁_h_, the methylation reaction propensities are calculated as

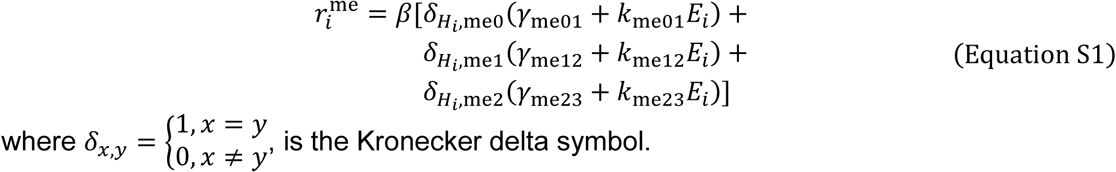

where 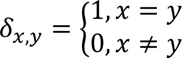, is the Kronecker delta symbol.

To incorporate the positive feedback between PRC2 and H3K27me3/me2 (but not H3K27me1), the model allows these two repressive marks to activate PRC2 and promote methylation on neighboring histones (*75*). Therefore, we defined an enhancement factor (dimensionless) for histone 𝑖, denoted as 𝐸_𝑖_ in Equation S1, to capture this effect

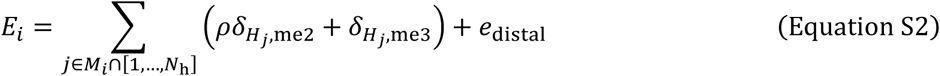

where 𝑀_𝑖_ is the neighboring histones, which is defined as

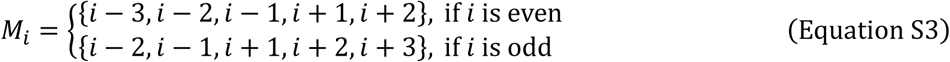

In our model, the first term in Equation S2 accounts for the proximal gain, and 𝜌 represents the reduced efficiency (∼10-fold) of activated PRC2 by H3K27me2 relative to H3K27me3 (*75*). The second term, 𝑒_distal_ in Equation S2 captures the contribution of long-range interactions, or distal gain, when DNA looping brings together nucleosomes that are far from each other in one dimension (*79*), although this event is less frequent than the others. Note that, in Equation S2, histones outside the *STM* locus are ignored. As a result, this introduces a slight bias toward the active state, especially of histones on boundary nucleosomes, as the boundary histones only undergo one-sided recruitment. However, this bias is expected to be inconsequential.

In this study, we identified that PRC2 can be recruited to *STM* locus by two classes of transcription factor, i.e., *RABBIT EARS* (*RBE*) and *BASIC PENTACYSTEINE* (*BPC*), in a sequence-specific manner. This provides a great example and explanation for the biological background of the relative local PRC2 activity 𝛽 used in previous models (*40*). In our model, 𝛽 in Equation S1 is understood to be controlled by the polycomb response elements (PREs) and the local activity of RBE/BPC.

###### H3K27 demethylation

In plants, there exist several demethylases, e.g., jumonji-C domain-containing (JMJD) protein family, that can catalyze H3K27 demethylation (*80*, *81*). Our model applies non-processive demethylation, with one methyl group removal each time. To simplify the model, we no longer considered JMJD alone in the simulation, instead we used a combined demethylation rate, 𝛾_dem_ as the integrated effect of JMJD-dependent and random removal of methyl groups (Fig. 4A and fig. S10A). Therefore, each histone 𝑖 undergoes demethylation with propensity

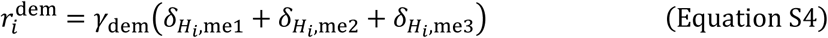

Furthermore, the model assumes that H3K27 demethylation is also coupled with gene expression event in two ways (*40*) (Fig. 4A and fig. S10A). On one hand, every transcription can result in random histones methylation removal (one methyl groups at a time) at each histone as the RNA polymerase moving (*44*), with probability denoted as 𝑃_dem_ per histone. Therefore, the process is shown as

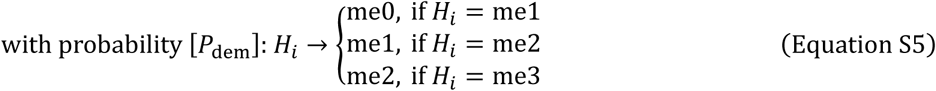

On the other hand, histone turnover is positively correlated with gene transcriptional activity (*45*, *82*), hence we refreshed each nucleosome (a pair of histones) at each time with exchange probability 𝑃_ex_ per histone independently. As a result, the probability that a nucleosome will not be replaced is the product of its two histones, i.e., (1 − 𝑃_ex_)^2^, therefore the turnover process is shown as

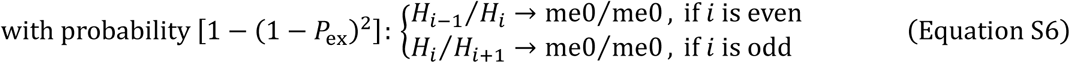

##### *STM* Expression Cycle

###### Gene transcription

*STM* expression, as an important component of our model, is linked to both epigenetic modification and stem cell division (Fig. 4A and fig. S10A). Meanwhile, STM protein itself also plays an important role in maintaining the undifferentiated state of pluripotent cells in shoot apical meristem (SAM) and/or axillary meristem (AM) (*83*, *84*). To investigate the repressive effect of epigenetic modification on gene transcription, we evaluated the dependence of mRNA production on H3K27me3/me2 levels. The transcriptional initiation rate, 𝑓 is a piecewise linear function of the average repression level across the *STM* locus, multiplied by another two trans-activation factors (discuss later). Altogether, the formula is given by

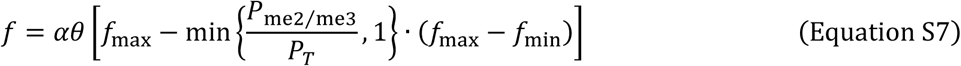

where 𝑓_max_ (𝑓_min_) is upper (lower) bound and 𝑃_me2⁄me3_ is the proportion of H3K27me3/me2

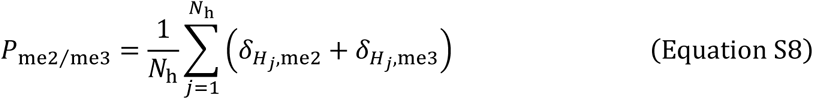

The threshold, 𝑃_𝑇_ in Equation S7 captures the sub-saturating level for H3K27me3/me2 to reach the maximal repression effect (*46*), the rationality of which was justified by fitting mass spectrometry data from stable isotope labeling by amino acids in cell culture (SILAC) (*40*, *85*). Moreover, previous model set an upper bound 𝑓_lim_ for the transcription initiation rate to restrict it in a reasonable range (*40*, *43*); however, we found that this cannot reproduce all of our experimental results. Instead, we successfully addressed this problem by adding a very low rate, 𝛾_transcr_ for noisy transcription. Finally, the propensity of *STM* gene transcription is

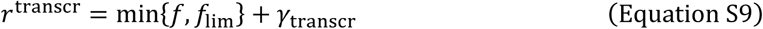

In *Arabidopsis*, a previous study revealed a self-sustained loop of *STM* gene in leaf axil to maintain its expression: STM interacts with ATH1, expressed by gene *ARABIDOPSIS THALIANA HOMEOBOX GENE1* (*ATH1*), to form a ATH1-STM complex, subsequently this complex binds to the *STM* locus by ATH1 and activates gene transcription (*8*). This process is reminiscent of a ligand-receptor interactions, hence we defined a self-activation factor for this path, denoted as 𝜃 in Equation S7, using the Hill equation

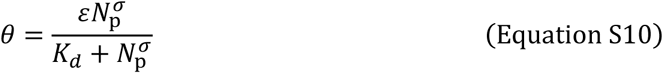

where 𝜀, 𝐾_𝑑_ and 𝜎 is the maximum self-activation level, apparent disassociation constant and Hill coefficient, respectively. Note that, 𝜀 is determined by the affinity between ATH1*-*STM complex and *STM* locus, thus this parameter actually only varies within a small range, i.e., hardly raises, due to the finite length of *STM* gene and steric effect of binding, although the number of ATH1 protein molecule can be infinite in the system (Fig. 5B). Moreover, we set 𝜎 > 1 to reflects the synergistic effect.

In our experiments, we manipulated the cell cycle by overexpressing KRP4/CYCD3;1, both of which could act on cyclin-dependent kinases (CDKs), the key-regulator of cell cycle. KRP4 is a CDK inhibitor (CKI), while CYCD3 can form active complex with CDK4/6 (*17*). However, CDK activity correlates with stem cell identity maintenance in meristem (*86*). Apart from physical dilution effect of epigenetic marks (Fig. 3G), in general, we considered two possible molecular pathways, one involving direct action on genes and one involving indirect action on chromatin modification. In our simulation, we took the direct way as major model, while indirect way as alternative model (see Additional Details).

For the direct way, we thought the observed correlation might be explained by *RETINOBLASTOMA-RELATED* (*RBR*), a negative regulator of cell cycle in plants that is inactivated by CDKs (*17*). Overexpressing RBR promotes meristem cell differentiation (*87*, *88*), while a reduction in RBR leads to stem cell accumulation and differentiation defect (*87*, *89*). Moreover, transient overexpression of *RBR* decrease the abundance of *Nicotiana tabacum homeobox 15* (*NTH15*, a *KNOX* factor, the ortholog of *Arabidopsis STM*) transcripts in tobacco SAM (*88*). During the autotrophic stage of *Arabidopsis*, RBR directly binds to promoters to silence specific genes (*90*), but whether *RBR* takes this mechanism or another pathway in meristem is still unclear. However, these results suggest that there do exist a complex regulatory network in meristem. To fit our experimental result and simplify the model, we used the simplest power function to represent the mathematical relationship (net effect, like a compound function) between the two outputs of the network, i.e., division frequency (inverse of cell cycle 𝑇 ) and cycle-dependent *STM* activation factor 𝛼

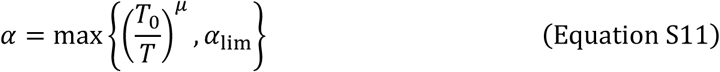

where 𝑇_0_ is the normalized cell cycle, 𝜇 is the exponential scaling factor, and 𝛼_lim_ is the lower bound, respectively. Besides, previous works suggested that the normal cell cycle in meristem typically ranged from 18-36 hours (*91*, *92*), and we fixed 𝑇_0_ to 22.0 hours to follow previous works (*40*, *43*). For biological continuity, if cell cycle changes in our simulations, we introduced a buffering period with a constant exponential changing rate 𝐹, therefore the real activation factor 𝛼 in Equation S7 is given by

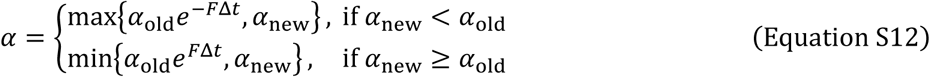

where 𝐹 is positive and 𝛼_old_ (𝛼_new_) is computed from Equation S11 using cell cycle before (after) change, respectively.

In the end, transcription events are coupled with another two events: demethylation and histone exchange, which we have already elaborated above in the section *H3K27 demethylation*.

###### mRNA translation

To simplify the model, we no longer considered the conversion of the *STM* gene to mRNA (transcription) and of the mRNA to STM protein (translation) as two split processes, instead we combined them into a single step, i.e., gene expression. The purpose of this approach is to eliminate an extra Gillespie’s simulation of intermediate state mRNA (*93*), which is actually not important in this model mechanism. However, to integrate transcription and translation, we introduced another parameter, 𝑛_ppt_ to denote the average protein molecule translated per mRNA transcript (where “ppt” stands for protein per transcription) (Fig. 4A and fig. S10A). Once a gene transcription event occurred in the simulation, we added the STM protein number by 𝑛_ppt_. Thus, the protein number in our model is more like a generalized concept, as 𝑛_ppt_ can also take float values. Here we ignored the time interval between each translation of the same mRNA transcript.

###### Protein degradation

In our model, we assumed that the degradation probability was constant. Therefore, the propensity for protein degradation is proportional to the remaining protein molecule number in the system, which is computed as

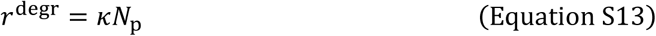

where 𝜅 is the degradation coefficient. Once a protein degradation event occurred in the simulation, we subtracted the number of STM protein molecule by 1. If the remaining molecule number is less than 1 before subtraction (possibly happen when 𝑛_ppt_ is not an integer), then we set it to 0 to avoid generating a negative value.

##### DNA Replication and Cell Division

In the simulation, DNA replication is inserted into the normal Gillespie process as a truncated event when time of the next coming event exceeds the next replication time (*94*). Actually, experiments suggested that H3/H4 tetramers did not dissociate and were shared evenly between daughter strand (*95*). Therefore, when replication is carried out, each nucleosome (a pair of histones) is updated with a unmethylated one with a probability of 0.5 (*40*). After each DNA replication, since we only tracked one daughter strand randomly

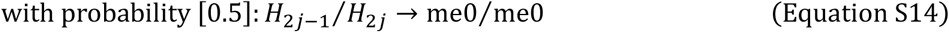

Moreover, we performed equal-scaling transformation when cell cycle changed in simulation, e.g., if a cell had 1.5 hours to the next division, and then cell cycle doubled, the transformed cell divided 3 hours later.

##### Additional Details

###### Gene activity

Defined as the average transcription number in a cell population per 15 minutes interval.

###### Cell division arrest

To simulate the situation when cells stop dividing by specific inductor, we allowed the cell cycle to be infinite in our simulations. If cell division is arrested, DNA replication events are excluded from the Gillespie process.

###### Protein stable/metastable point

According to our model, the propensity for *STM* protein degradation is linear, while the propensity of STM protein production (gene expression) is sigmoid and fluctuates between the two extreme epigenetic state (fig. S10B). For a given situation, the intersections of these two curves represent the fixation points of protein in the ideal system. Regardless, there should be one fixation point (≈ 0) due to the noisy transcription, which stands for the stably differentiated state (fig. S12, solid blue line). However, the other two fixation points (if both exist) have very different properties. For the larger one, the system reaches a stable equilibrium and tends to recover itself like a spring after perturbation in any direction; thus, we termed this a stable point. Actually, this is the case for normal pluripotent cells with stable *STM* expression (fig. S12, solid red line). For the smaller one, a positive disturbance leads to continuous increase until stable point, which represents the direction of pluripotency restoration, while a negative disturbance causes irreversible decrease until vanish and maintain itself from then on, which represents the direction of differentiation. Altogether, the equilibrium here is unstable and we named it as metastable point, or critical point (fig. S12, green dashed line). When methylation level rises, the stable point drops while the metastable point elevates until they meet at branch point (saddle point) and annihilate, which is similar to saddle-node bifurcation process in mathematics.

###### Differentiation/pluripotency criteria

We used the remaining number of STM protein molecule in the system to characterize the cell fate. In our simulation, the cells with no STM protein left were viewed as completely differentiated, while the cells with STM protein number more than half of the stable point in the lowest epigenetic repression (H3K27me0) state were viewed as stem cells, others are considered in a transitional state.

###### Bistability measurement

Bistability (fig. S10, C to E) describes the ability to maintain both stem cell fate with low histone repression (active transcription) and differentiated cell fate with high histone repression (inactivate transcription) state of *STM* gene. Here we followed the conception of previous models in which the bistability measurement (*40*, *43*, *96*), 𝐵 was given by

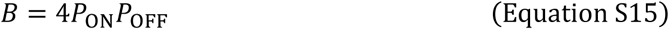

where 𝑃_ON_ stands for the probability of the *STM* gene being in the epigenetic ON state, which is defined as

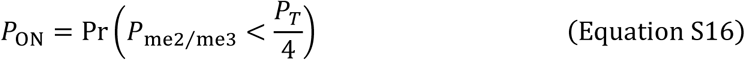

and likewise, 𝑃_OFF_ represents the probability of the *STM* gene being in the epigenetic OFF state

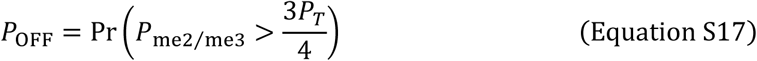

In practice, 𝐵 is calculated using Equation S14 from the combined outputs from the equal number of simulations initiated from either stem cells (uniform H3K27me0, an STM protein level at high stable point) or differentiated cell (uniform H3K27me3, no STM protein). 𝐵 values closer to 1 indicate stronger bistability. For reliability, we allowed the system to recover from the perturbation of DNA replication before sampling, thus only the data from stable state (last hour of each cell cycle) was used (*40*).

###### Alternative model

As mentioned above, there is another possible pathway in which cell division frequency coupled with histone methylation indirectly affects *STM* transcription. Although the underlying molecular mechanism in plants remains unclear, some studies have suggested that CDK1 could phosphorylate the catalytic subunit of PRC2 at specific target and disrupted its activity in human cells (*97*). Therefore, we also proposed an alternative model in which an increase in cell cycle promote PRC2 activity instead of blocking gene transcription directly (fig. S13). In this case, the propensity of histone methylation (Equation S1) and *STM* transcription (Equation S12) are replaced in respective with

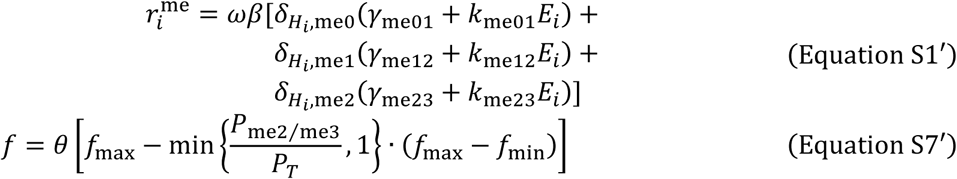

In equation S1′, the cycle-dependent PRC2 activation factor 𝜔 is given by simple sigmoid function

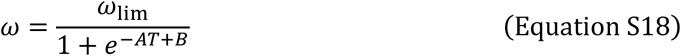

where 𝑇 is cell cycle, 𝐴 is the absolute of slope, 𝐵 is the intercept and 𝜔_lim_ is the upper bound. Here we omitted the buffering period of 𝜔.

Note that, the net effect of both the major/alternative model is decreasing the *STM* transcription efficiency. Thus, as expected, the alternative model gave very similar trajectories of tendency to major model, but with a slower silencing velocity, which is consistent with its indirect manner (Fig. 4, B and C, fig. S11, A to H and fig. S12, A to H).

###### Cycle-dependent activation

In major/alternative model, the role of cycle-dependent activation factors, i.e., 𝛼 and 𝜔, is to alter the fixation point (corresponds to stable state) in system phase space. The dilution of H3K27me3 when DNA replication, in theory, can only change the initial position in the next cycle but not the structure of dynamic system.

##### Programming Languages and Code Resources

The model programs were written in MATLAB R2024a and Python 3.12. All simulations were tested both on the Windows (win10/win11) and Linux (Ubuntu 20.04/22.04 LTS) operating system. The source code of the model is available at https://github.com/yy420106/stmExpr-chromMod-cellDivModel.

---

**Fig. S1.**
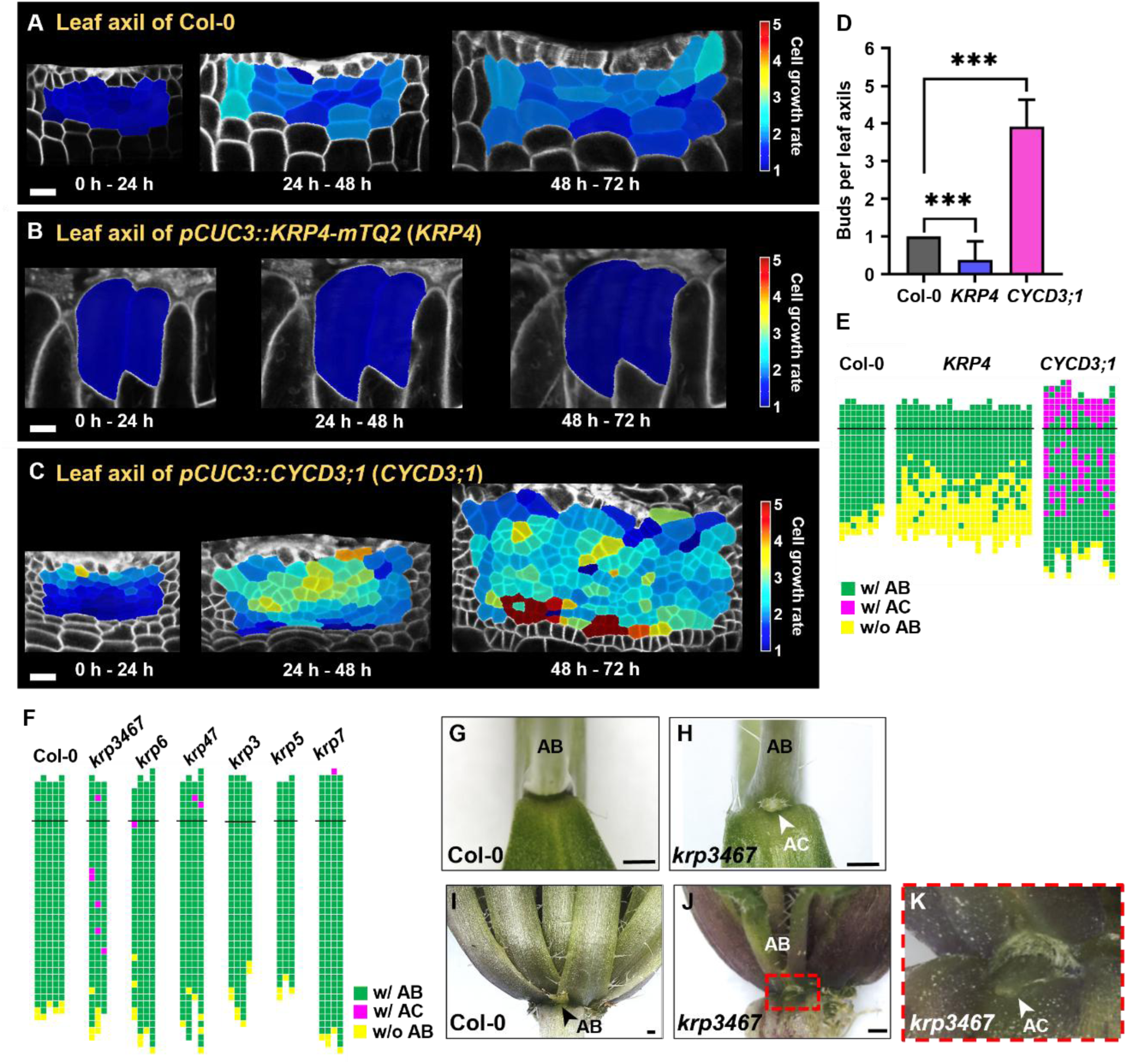
Changes in the rate of cell division affect AM initiation. (**A‒C**) Heatmaps showing the leaf axil cell growth rates of Col-0 (A), *KRP4* (B) and *CYCD3;1* (C) plants. Scale bars, 20 µm. (**D**) Statistical analysis of the number of axillary buds in detached Col-0 (Fig.1K), *KRP4* (Fig. 1L) and *CYCD3;1* (Fig. 1M) leaves. The error bars indicate the standard deviation (SD). ***p < 0.001 (Student’s *t* test). (**E**) Schematic diagrams of axillary bud formation in Col-0, *KRP4*, and *CYCD3;1* plants. Each column represents a single plant, and each square represents a leaf axil. The thick black horizontal line represents the border between the youngest rosette leaf and the oldest cauline leaf. The bottom row represents the oldest rosette leaf axils, with progressively younger leaves above. Green indicates the presence of a single axillary bud (w/ AB), yellow indicates the absence of an axillary bud (w/o AB), and magenta indicates the formation of an extra accessory bud (w/ AC). (**F**) Schematic diagrams of axillary bud formation in Col-0 and *krp3467*, *krp6*, *krp47*, *krp3*, *krp5*, and *krp7* mutants. The diagram components are as described in (E). (**G‒K**) Close-up of cauline leaf (G and H) and rosette leaf (I‒K) axils in Col-0 (G, I) and *krp3467* (H, J, K), showing the presence of axillary buds or branches (ABs) and accessory buds (ACs). Note that (K) is an enlargement of (J). Scale bar, 500 µm.

**Fig. S2.**
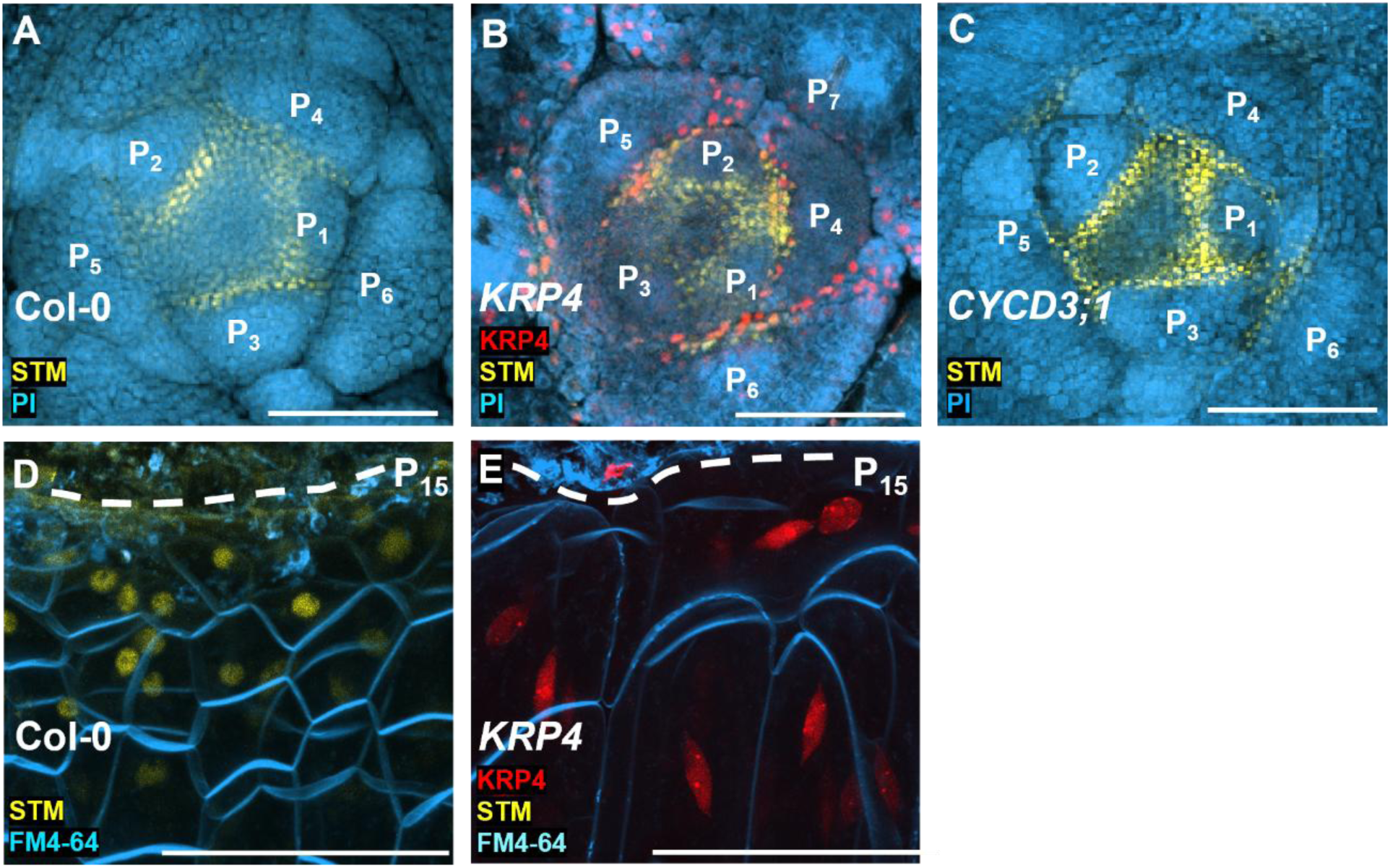
Changes in the rate of cell division affect *STM* expression. (**A‒C**) Transverse sections of the vegetative shoot apices of Col-0 (A), *KRP4* (B) and *CYCD3;1* (C) plants stained with PI (cyan) and showing *STM* (yellow) and *KRP4* (red) expression (B). Note the reduced *STM* signal in P_7_ and the older leaf axils. Scale bars, 50 µm. (**D** and **E**) Leaf axil *STM* signals (yellow) in Col-0 (D) and *pCUC3::KRP4*-*mTQ2* (*KRP4*, E) in P_15_, with FM4-64 staining (cyan). Red indicates *KRP4* expression (E). Note the lack of *STM* signals in *KRP4*. Scale bars, 50 µm.

**Fig. S3.**
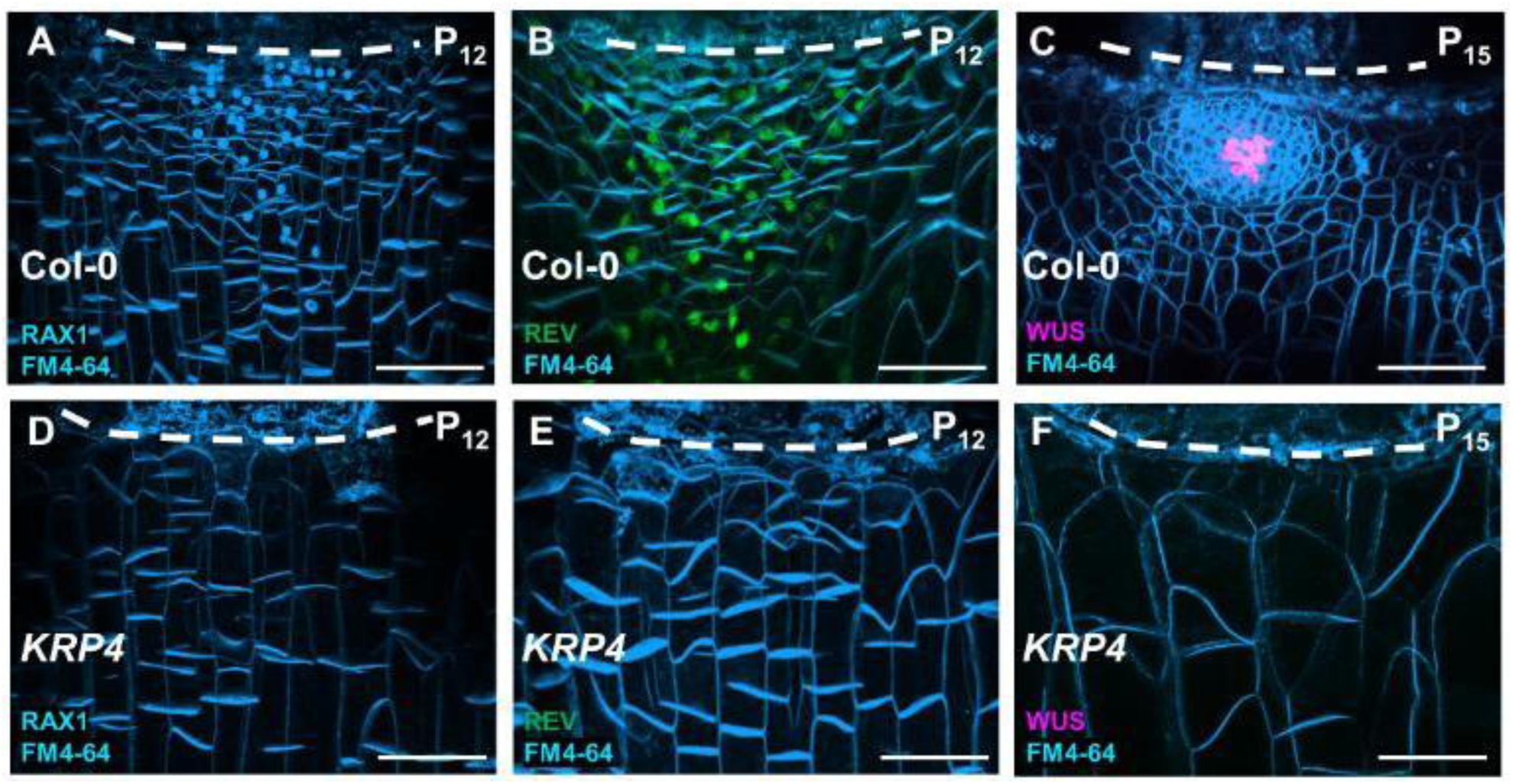
AM initiation-related genes were inhibited in the axils of *KRP4.* Expression of *RAX1* (cyan), *REV* (green), and *WUS* (magenta) genes in the leaf axils of Col-0 (**A‒C**) and *KRP4* (**D‒F**) with FM4-64 staining (cyan). The white dotted lines indicate the boundary of the leaf axil. Scale bars, 50 µm.

**Fig. S4.**
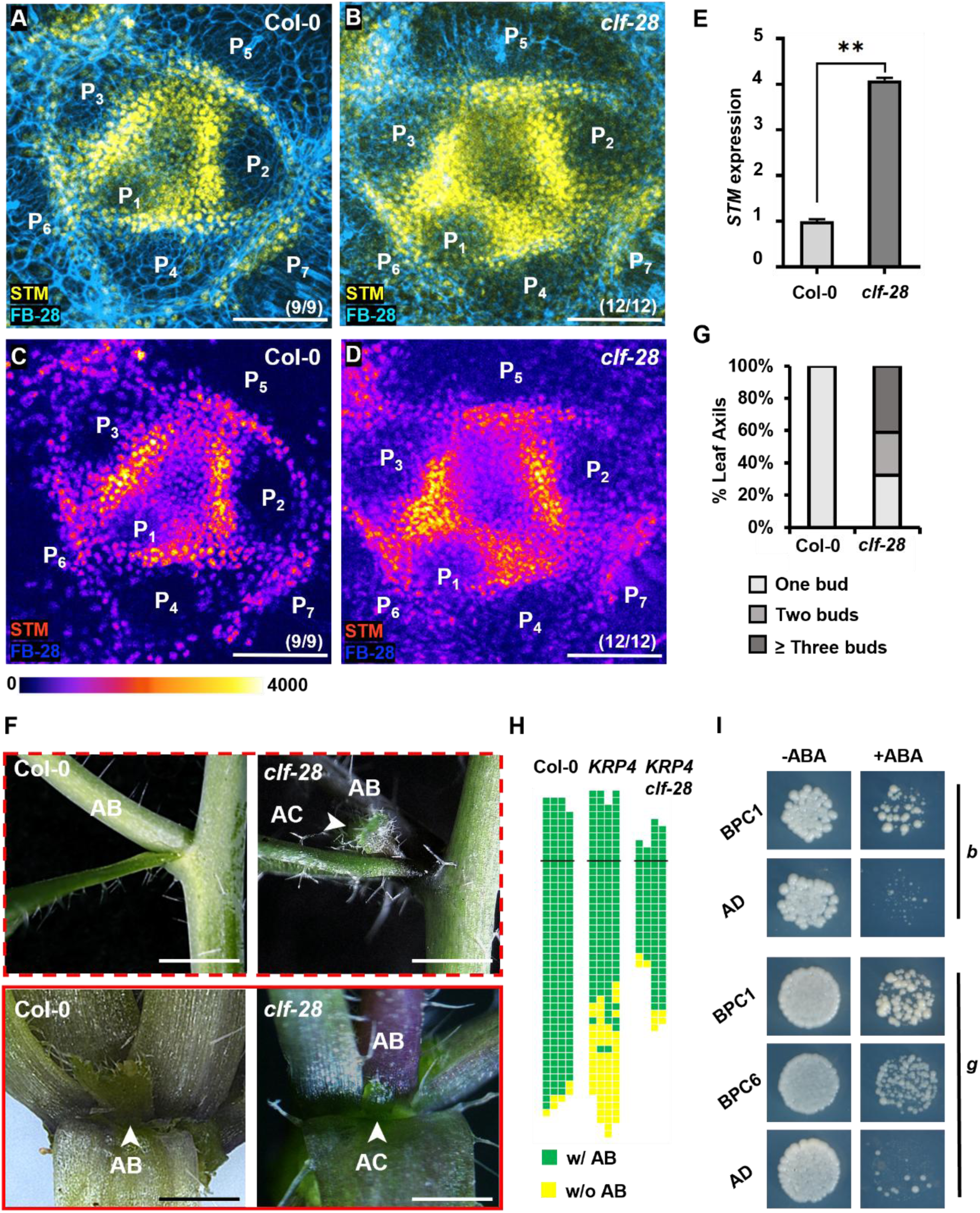
*STM* expression in leaf axils was increased in *clf-28*. (**A‒D**) Transverse sections of vegetative shoot apices of Col-0 (A) and *clf-28* (B) plants showing STM (yellow) expression with FB28 staining (cyan). The two images below the transverse sections show the corresponding heatmaps (C and D). (m/n) indicates that m in n biological repeats shows the displayed features. Scale bars, 100 µm. (**E**) RT-qPCR analysis of *STM* expression in Col-0 and *clf-28* vegetative shoot apex tissues (with leaves removed) after 14 d under short-day growth conditions. The vertical axis indicates the relative mRNA transcript levels. The error bars indicate the SDs of three biological replicates performed in triplicate. **p < 0.01 (Student’s *t* test). (**F**) Close-up of the leaf axils in Col-0 and *clf-28*. Note, rosette leaf (red solid boxes) and cauline leaf axils (red dotted boxes) for Col-0 and *clf-28*, showing the presence of accessory buds (ACs, arrows) in *clf-28*. Scale bars, 1 cm. (**G**) Statistical analysis of axillary bud development in P_12-15_ leaf axils detached from Col-0 (F) and *clf-28* (G) plants. (**H**) Schematic diagrams of axillary bud formation in Col-0, *KRP4*, and *KRP4 clf-28* plants. The diagram components are as described in Fig. 2D. (**I**) Y1H assay showing the interaction of BPC1 and BPC6 with the *STM* locus fragments, as shown in Fig. 2A. The transcription factors BPC1 and BPC6 were fused to the GAL4 activation domain (AD). AD alone served as a negative control.

**Fig. S5.**
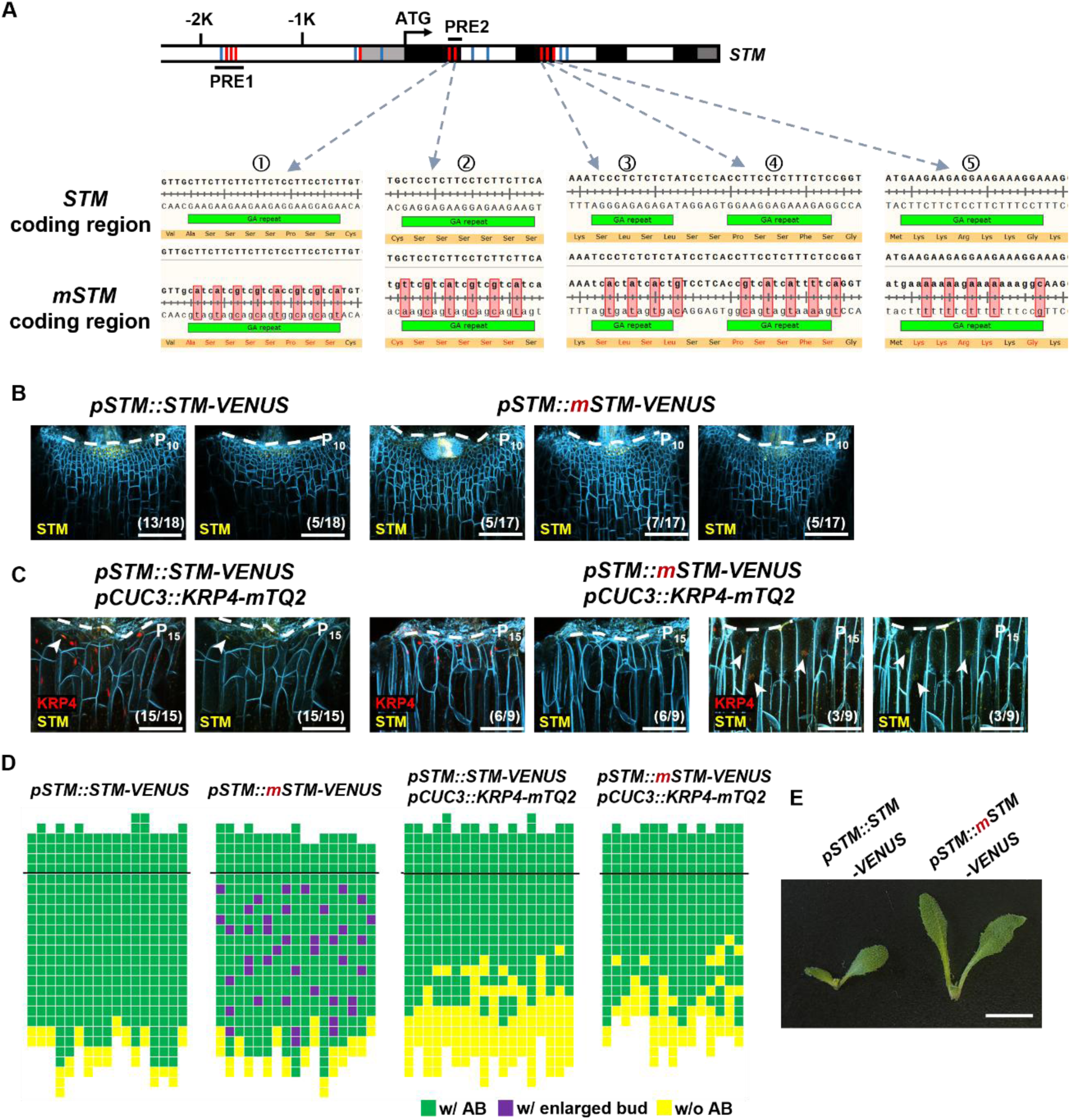
Synonymous mutations in the PRE element of the STM coding region cause elevated *STM* expression and increased cell division. (**A**) Synonymous mutations in the PRE element of the STM coding sequence. Parts 1 to 5 respectively show the five GA repeats located in the first and second exons of the *STM* gene. The first row displays the nucleotide and amino acid sequences of the GA repeat in the *STM* CDS, while the second row presents the corresponding sequences after mutation. The green horizontal bar indicates the position of the GA repeat sequence. (**B**) *STM* expression in the leaf axils of P_10_ was examined in the *pSTM::STM-Venus* and *pSTM::mSTM-Venus* (Synonymous substitution in the PRE element of the *STM* CDS) transgenic plants. The white dotted lines indicate the boundary of the leaf axil. Scale bar, 100 µm. (**C**) *STM* expression in the leaf axils of P_15_ was examined in the *pSTM::STM-Venus pCUC3::KRP4-mTQ2* and *pSTM::mSTM-Venus pCUC3::KRP4-mTQ2* plants. The white dotted lines indicate the boundary of the leaf axil. White arrows indicate the *STM*-expression nuclei. Scale bar, 100 µm. (**D**) Schematic diagrams of axillary bud formation in *pSTM::STM-Venus*, *pSTM::mSTM-Venus*, *pSTM::STM-Venus pCUC3::KRP4-mTQ2* and *pSTM::mSTM-Venus pCUC3::KRP4-mTQ2* plants. Purple indicates the presence of an enlarged axillary bud. See Fig. 2D for additional details. (**E**) Abnormally enlarged axillary buds were frequently observed in the *pSTM::mSTM-Venus* plants. Scale bar, 1 cm.

**Fig. S6.**
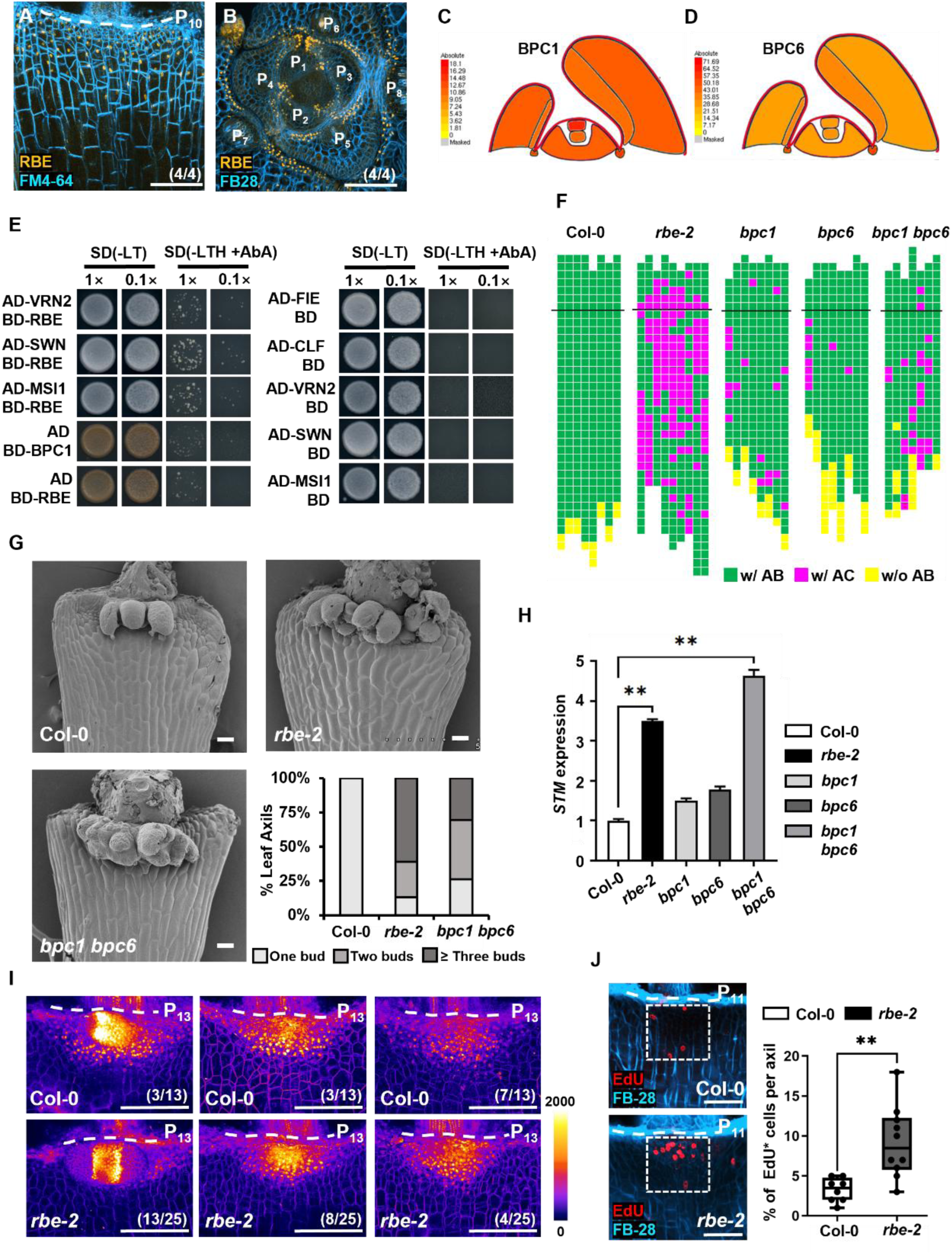
*RBE*, *BPC1* and *BPC6* inhibit AM initiation. (**A**) Expression of *RBE* (orange) in a leaf axil with FM4-64 staining (cyan) in *pRBE::GFP-RBE*. (m/n) indicates that m in n biological repeats shows the displayed features. Scale bars, 100 µm. (**B**) A transverse section of a vegetative shoot apex of a *pRBE::GFP-RBE* plant showing RBE (orange) expression, with FB28 (cyan) staining. (m/n) indicates that m in n biological repeats shows the displayed features. Scale bars, 100 µm. (**C‒D**) eFP browser showing the expression of the *BPC1* and *BPC6* genes in the shoot apex. (**E**) Validation of VRN2, SWN and MSI1 interactions with the RBE protein via Y2H assays. See Fig. 2C for additional details. (**F**) Schematic diagrams of axillary bud formation in Col-0, *rbe-2*, *bpc1*, *bpc6* and *bpc1 bpc6* mutants. See Fig. 2D for additional details. (**G**) Axillary buds in detached P_12-15_ leaf axils from Col-0 (upper left), *rbe-2* (upper right), and *bpc1 bpc6* (lower left) after 5 d under short-day growth conditions. Leaves were detached from plants grown under short-day conditions for 14 d. The lower right graph shows the ratios of axillary bud initiation of P_12-15_ in Col-0, *rbe-2* and *bpc1 bpc6*. Scale bars, 100 µm. (**H**) RT-qPCR analysis of *STM* expression in Col-0, *rbe-2*, *bpc1*, *bpc6*, and *bpc1 bpc6* vegetative shoot apices (with leaves removed) after 14 d under short-day growth conditions. The vertical axis indicates the relative mRNA transcript levels. The error bars indicate the SDs of three biological replicates performed in triplicate. *p < 0.01 (Student’s *t* test). (**I**) Heatmap showing *STM* expression in the leaf axil of P_13_ of Col-0 and *rbe-2*. Note the enhanced signal in *rbe-2,* especially in older leaves. (m/n) indicates that m in n biological repeats shows the displayed features. Scale bars, 100 µm. (**J**) EdU staining (6 h) of the leaf axil of P_11_ Col-0 (upper left) and *rbe-2* (lower left) plants. The box plot shows the number of nuclei labeled by EdU in each leaf axil in Col-0 and *rbe-2* (right column). Scale bars, 50 µm.

**Fig. S7.**
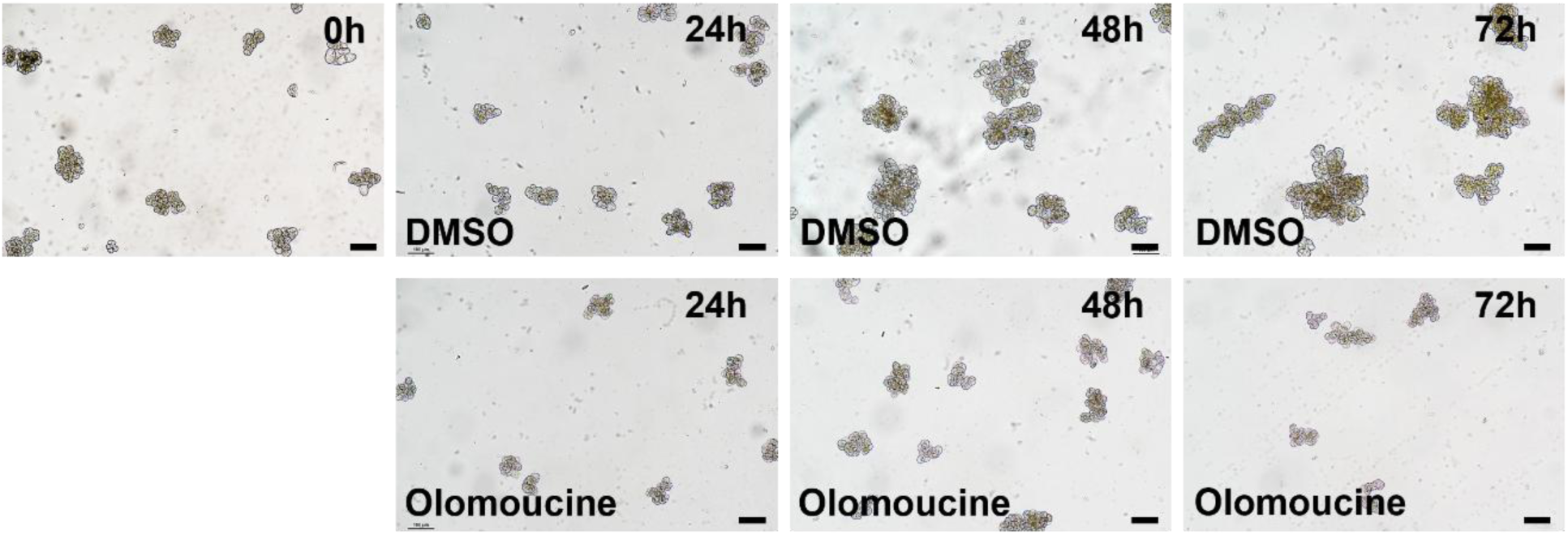
Cell cycle inhibitor arrested the cell division of T87 cells. Time-lapse imaging of T87 cell division after treatment with DMSO or olomoucine for 0h, 24h, 48h, and 72h. Scale bar, 100 µm.

**Fig. S8.**
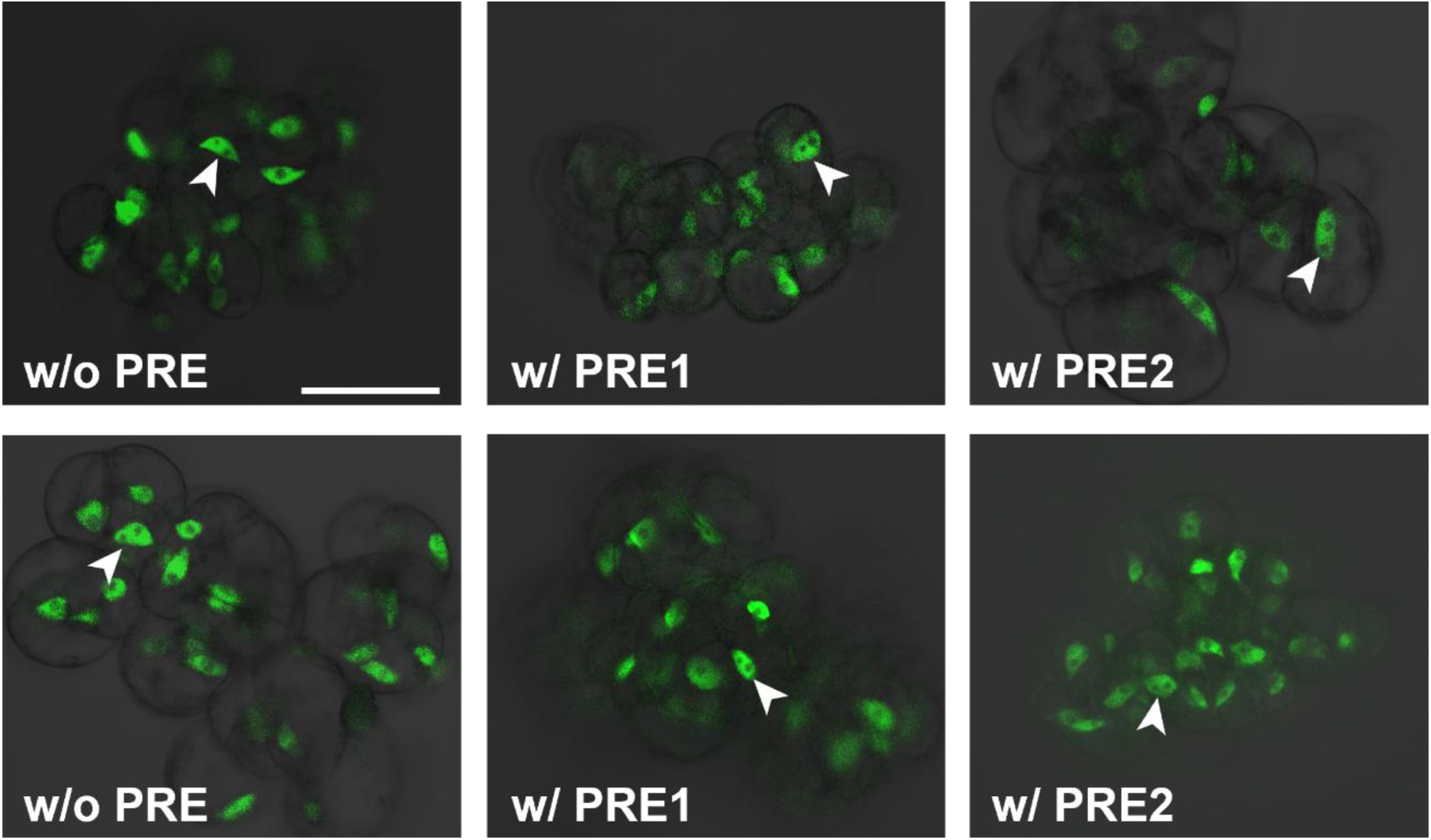
The control group (without the cell cycle inhibitor) showed active cell division in the cell mass. Both types of cell mass (with or without the PRE element) in the DMSO control group showed frequent cell division. Note, white arrows indicate the dividing nuclei. Scale bar, 50 µm.

**Fig. S9.**
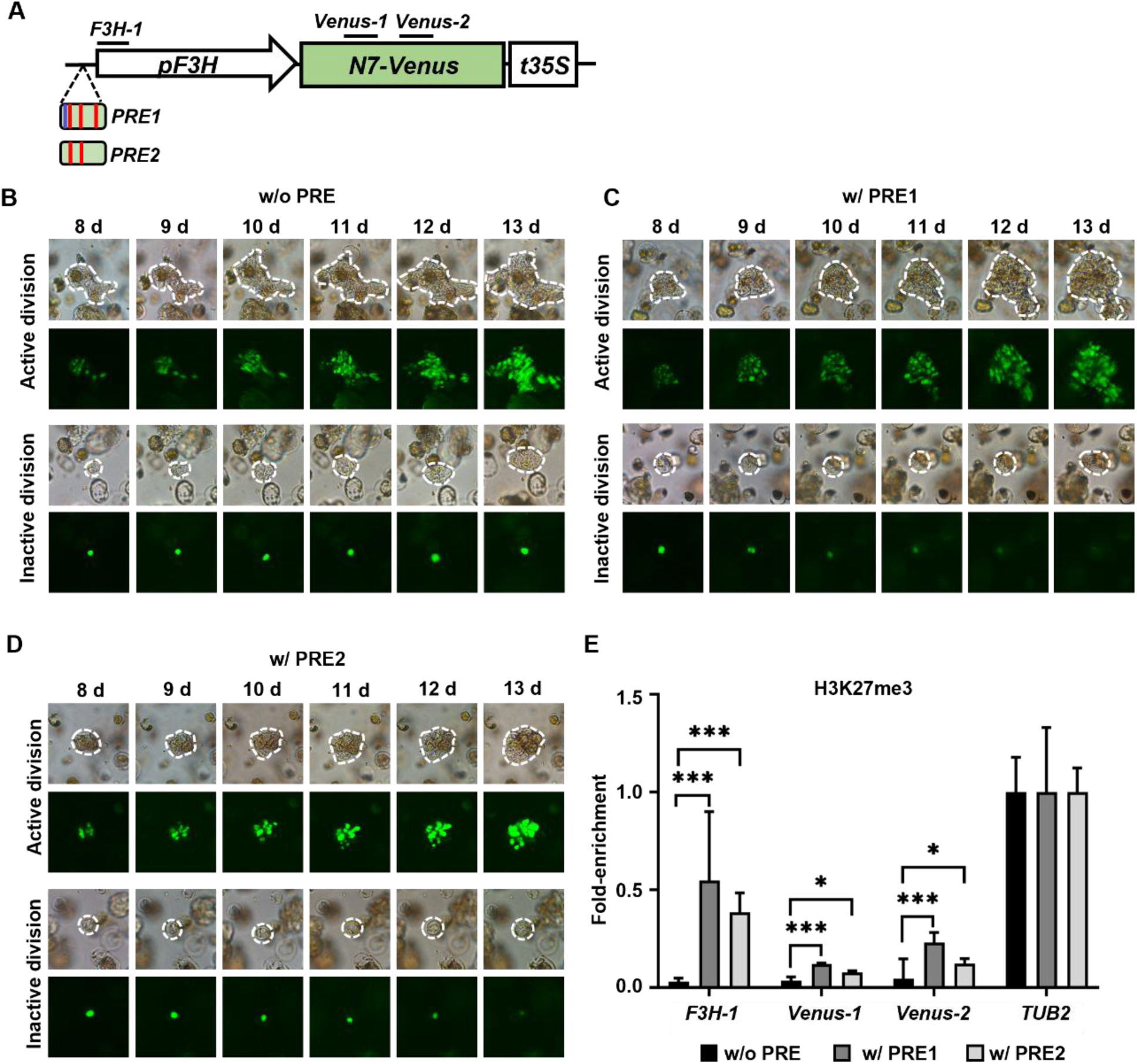
Cell division eliminates H3K27me3 deposition. (**A**) Schematic diagram of the constructs used to test the relationships among cell division, H3K27me3 modification, and gene expression. See Fig. 3A for additional details. (**B-D**) *Venus* expression dynamics in different cell division states in *pF3H::Venus* without (B) and after the addition of the PRE1 (C) or PRE2 (D) element after 8 d of resistance screening. Cells with active and inactive divisions are shown. The white dotted lines indicate the cells or cell clusters that were tracked. See Fig. 3, B and D for additional details. (**E**) ChIP-qPCR results showing a comparison of the effects of PRE insertion on H3K27me3 accumulation. Three fragments, *F3H-1*, *Venus-1* and *Venus-2*, which are indicated by the short lines in (A), were tested. The error bars indicate the SDs of three biological replicates performed in triplicate. *p < 0.05, ***p < 0.001 (Student’s *t* test).

**Fig. S10.**
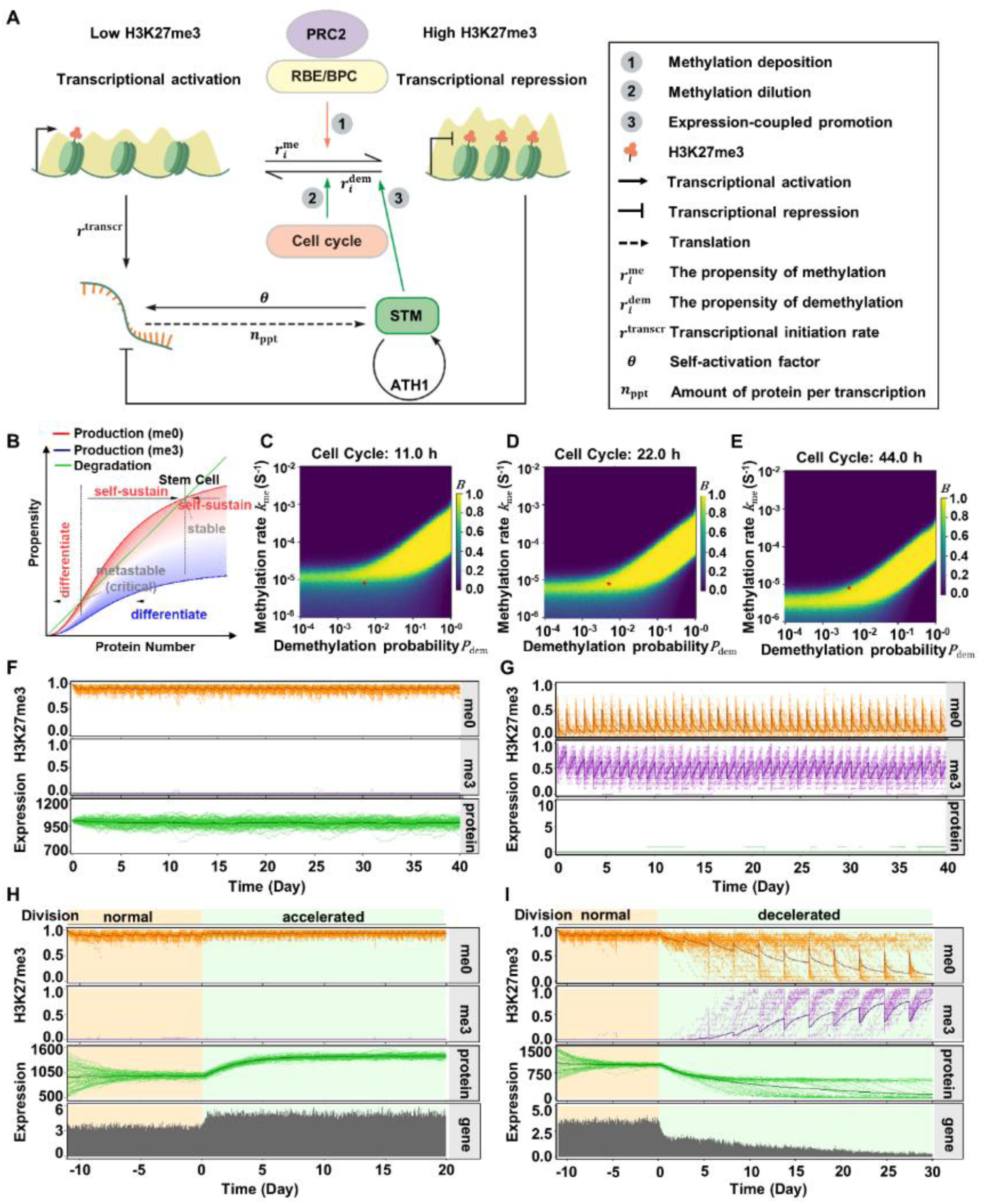
The hybrid histone modification-*STM* expression-cell division model. (**A**) Schematic of the model description showing the key feedback mechanisms. The basic diagram components are the same as those described in Fig. 6A. Additional model parameters have been added. **(B)** Main mathematical principles of the model. The shaded area represents the variation range of the *STM* expression curves between two extreme boundaries (H3K27me0/me3), with a redder/bluer curve representing a lower/higher epigenetic repression state. Note that the graph was not drawn to scale. (**C‒E**) Heatmap showing values for the bistability measurement 𝐵 , calculated from simulations. Each graph shows 𝐵 as a function of the methylation rate 𝑘_me_ and demethylation probability 𝑃_dem_ for the cell cycle at (C) 11 h, (D) 22 h, and (E) 44 h, as indicated at the top. 𝐵 is close to 1 for bistable systems. Red asterisks indicate the coordinates of the default values used in the model. For each parameter set, 100 simulations were initialized from either pluripotent cells (uniform H3K27me0, an *STM* protein at a high stable point) or differentiated cells (uniform H3K27me3, no *STM* protein) and simulated for 40 cell cycles. The results are averaged over all the simulations. The other parameters are fixed to their default values, as shown in Table S1. (**F** and **G**) Representative trajectories from stochastic simulations of (F) pluripotent cells (uniform H3K27me0, an *STM* protein at a highly stable point) and (G) differentiated cells (uniform H3K27me3, no *STM* protein). Each graph shows 80 overplotted trajectories. The thick line in each panel represents the average curve of the cell population. The values used in the simulation are shown in Table S1. (**H** and **I**) Representative trajectories from stochastic simulations of (H) 2-fold accelerated cell division and (I) decelerated cell division to 1/3 of the original rate. Simulations were started randomly from ±50% fluctuation of protein stable points in pluripotent cells (uniform H3K27me0) with a normal cell cycle (22 h), and 12 cell cycles were performed for preequilibration before changing the cell cycle. Each graph shows 80 overplotted trajectories. The thick line in each panel represents the average curve of the cell population. Gene activity was measured as the average number of transcription events in the cell population per 15 min interval. The other parameters are fixed to their default values, as shown in Table S1.

**Fig. S11.**
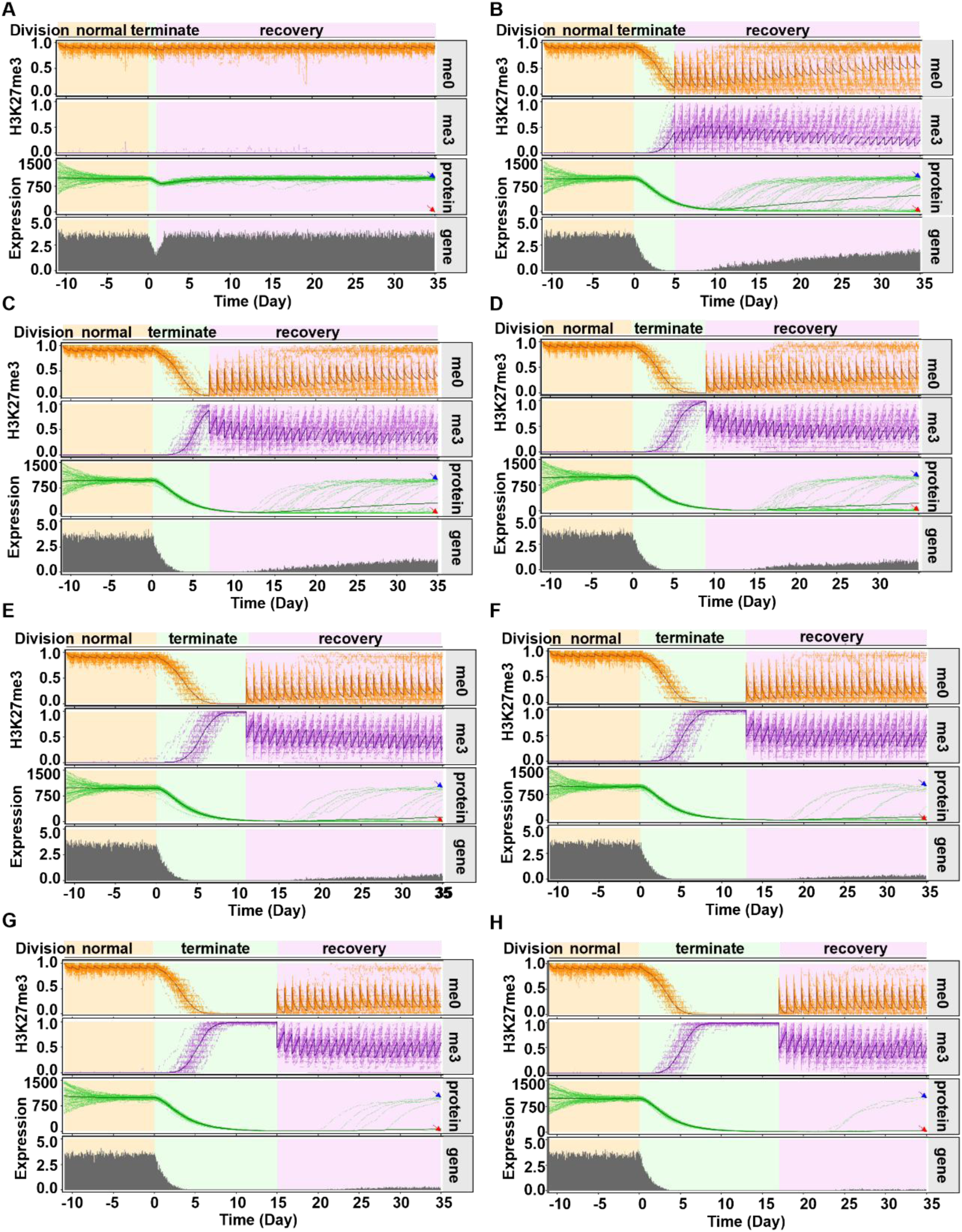
Additional justification and prediction of the hybrid histone modification-*STM* expression-cell division model. (**A‒H**) Representative trajectories from stochastic simulations of (A) 1 d, (B) 5 d, (C) 7 d, (D) 9 d, (E) 11 d, (E) 13 d, (G) 15 d and (H) 17 d of cell division arrest treatment before recovery. Simulations were started randomly from ±50% fluctuation of protein stable points in pluripotent cells (uniform H3K27me0) with a normal cell cycle (22 h), and 12 cell cycles were performed for preequilibration before quiescence initiation. Each graph shows 80 overplotted trajectories. The thick line in each panel represents the average curve of the cell population. Red arrowhead, differentiated group; blue arrowhead, pluripotency-restored group. Gene activity was measured as the average number of transcription events in the cell population per 15 min interval. The other parameters are fixed to their default values, as shown in Table S1.

**Fig. S12.**
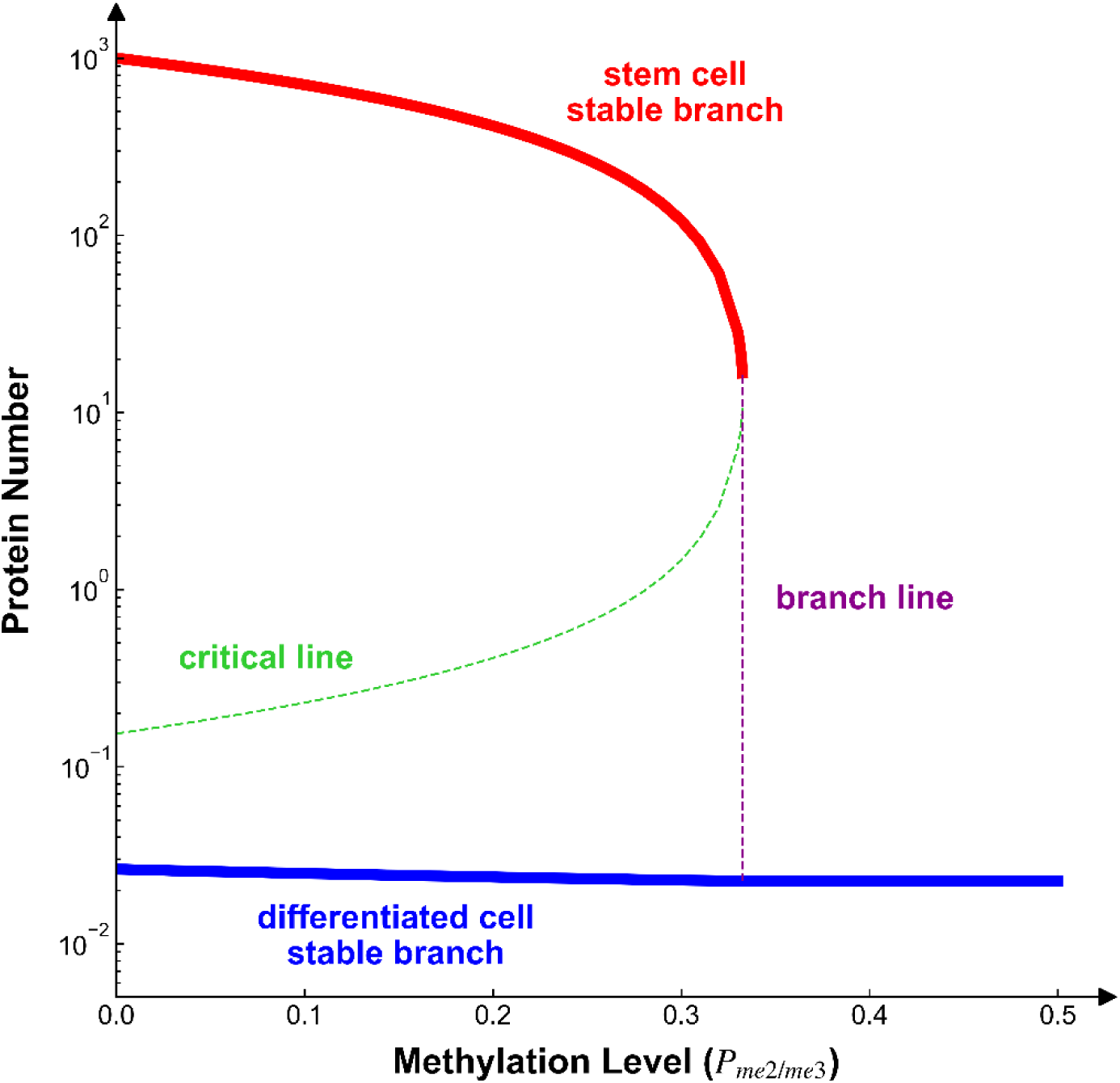
Cell fate bifurcation in model phase space. Solid blue line represents the stable branch of fully differentiated cells. Solid red line represents the stable branch of stem cells, that annihilates at the left limit of branch line (purple dashed line). Green dashed line represents the trajectory of critical/metastable point, which divide the left space into pluripotency bias and differentiation bias region.

**Fig. S13.**
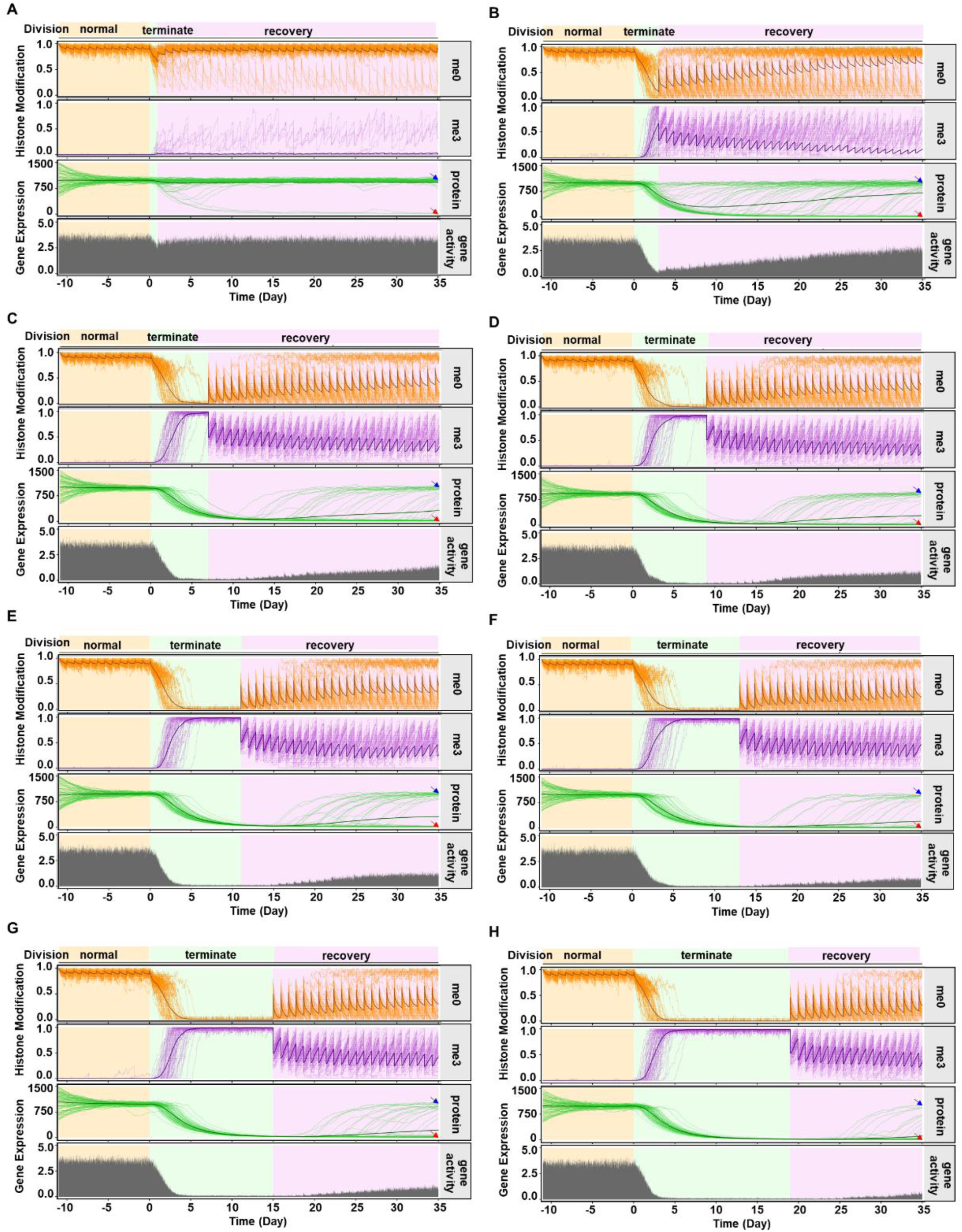
Prediction of alternative model. (**A-H**) Representative trajectories from stochastic simulations of (A) 1 day, (B) 3 days, (C) 7 days, (D) 9 days, (E) 11 days, (F) 13 days, (G) 15 days and (H) 19 days cell division arrest treatment before recovery using the alternative model, in which cell division frequency affects PRC2 activity instead. Simulations were started randomly from ±50% fluctuation of protein stable points in stem cell (uniform H3K27me0) with normal cell cycle (22.0 h), and performed 12 cell cycle for pre-equilibrium before quiescence. Each graph shows 80 overplotted trajectories. Histone modification level averaged within *STM* locus. Heavy line in each panel represents the average curve of the cell population. Red arrowhead, differentiated group; blue arrowhead, stem-restored group. Gene activity measured as the average number of transcription events in cell population per 15 min interval. Other parameters are fixed to their default values, as listed in Table S1. See also Fig. 4. B and C, and fig. S11, A to H.

**Table S1.**
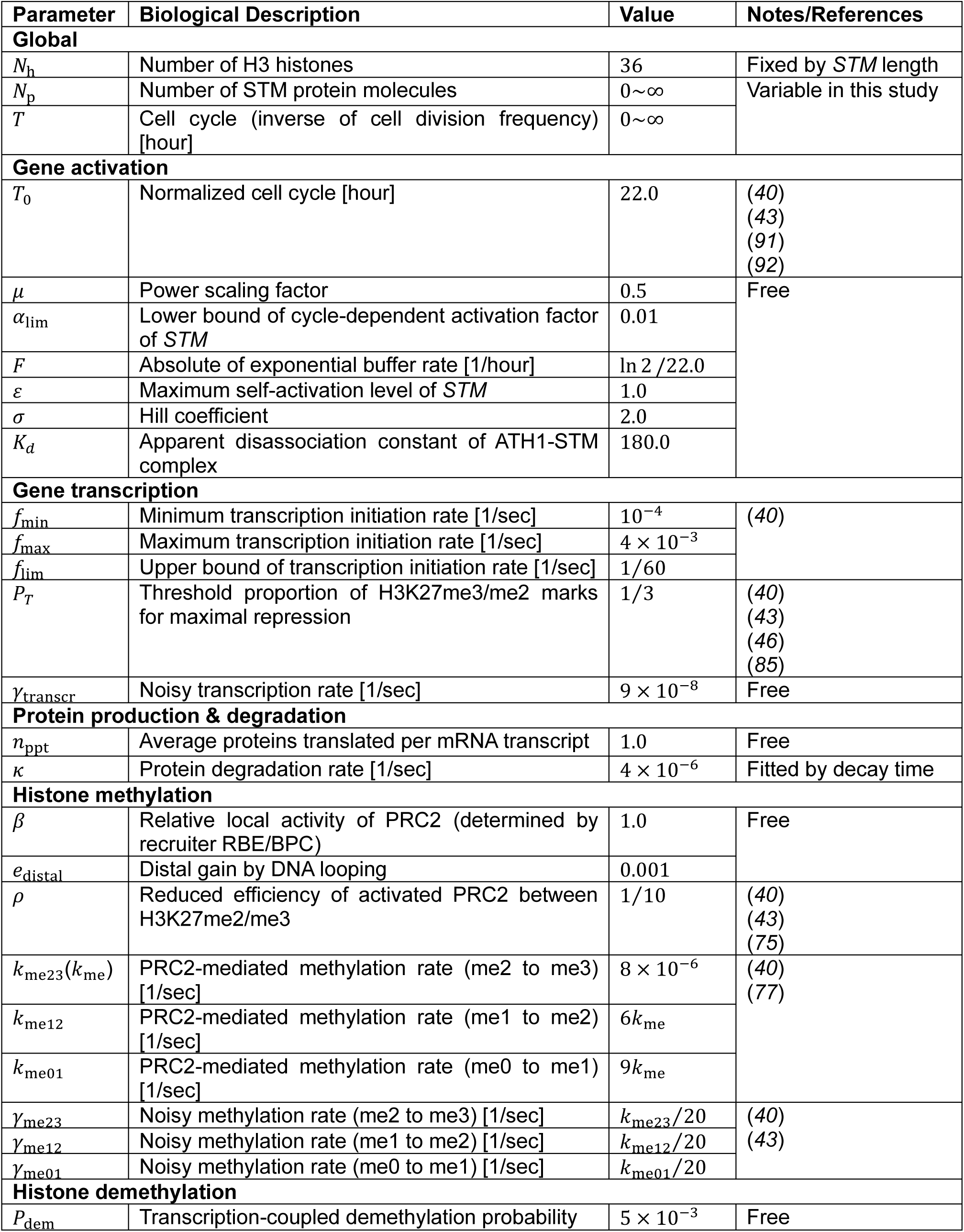
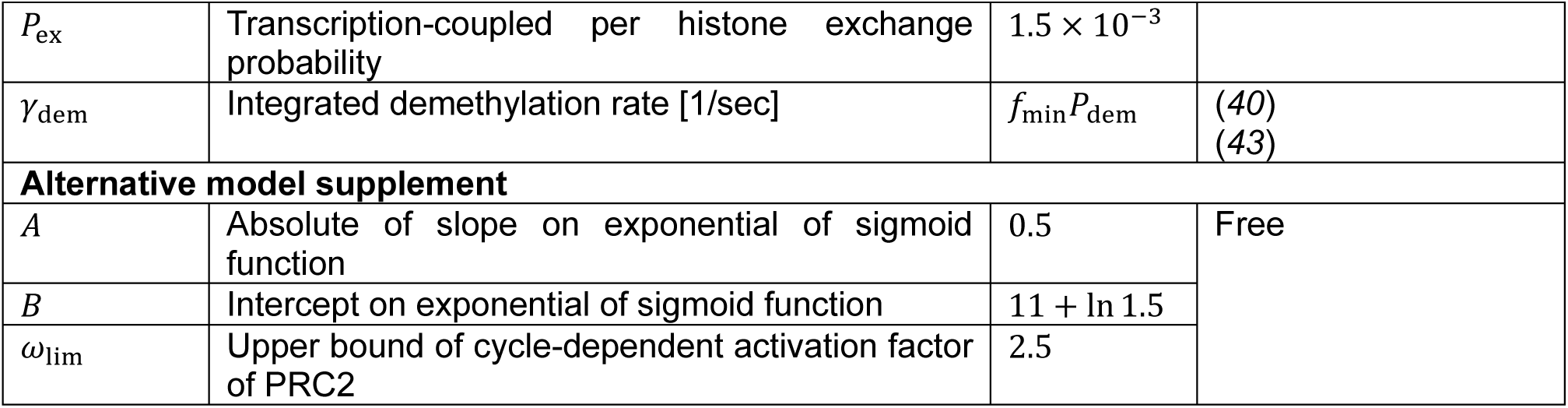
Summary of model parameters.

**Table S2.**
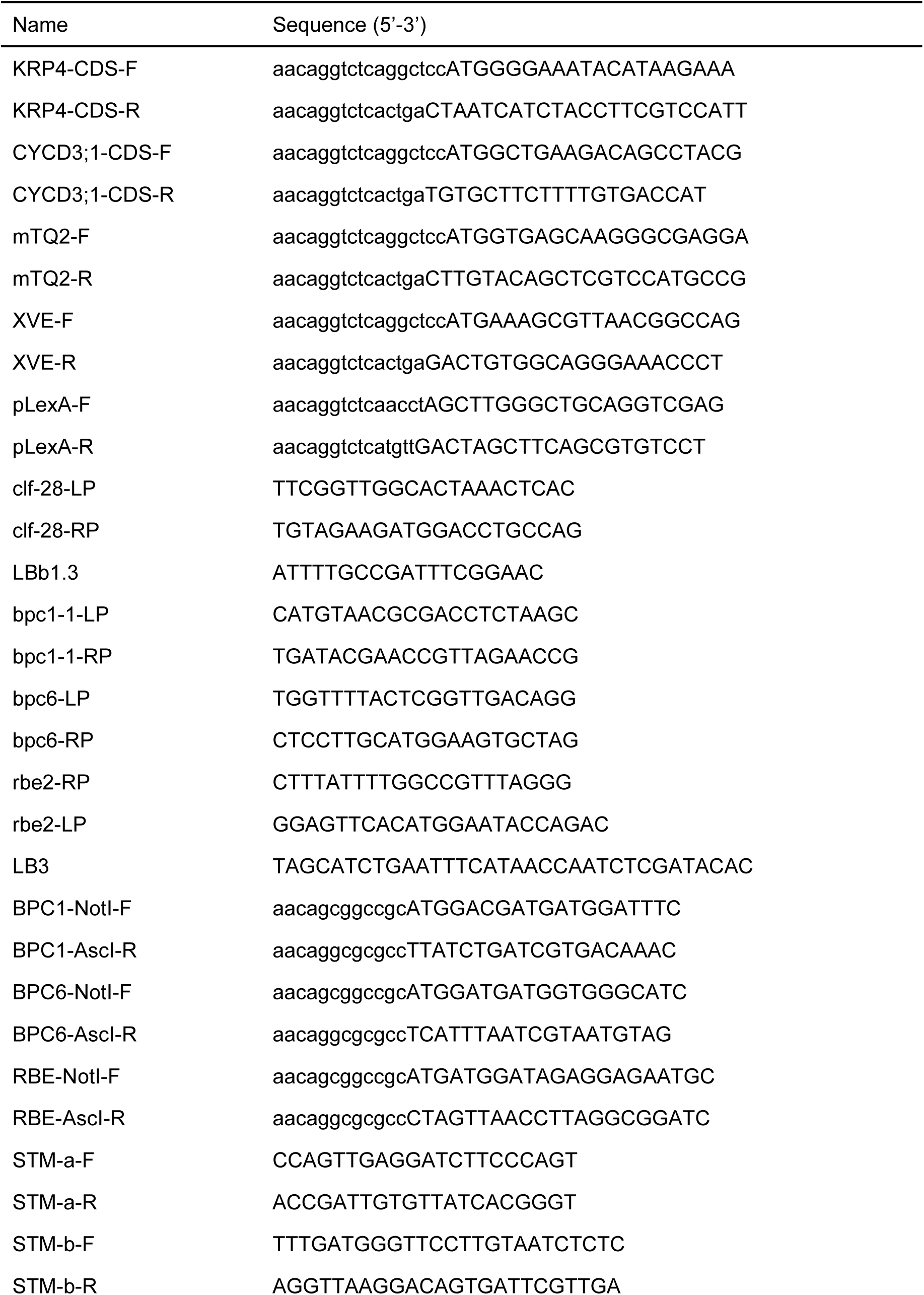
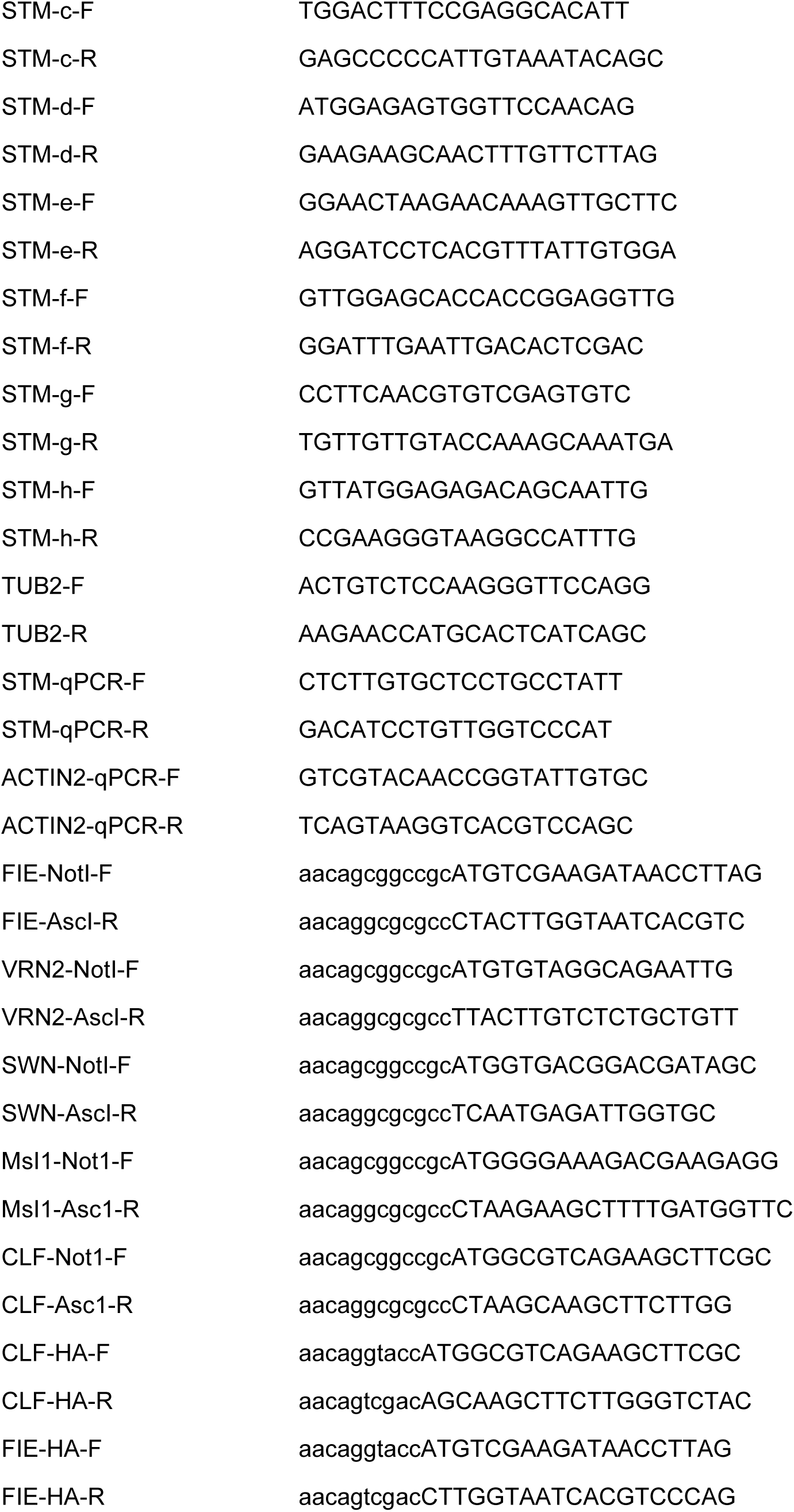
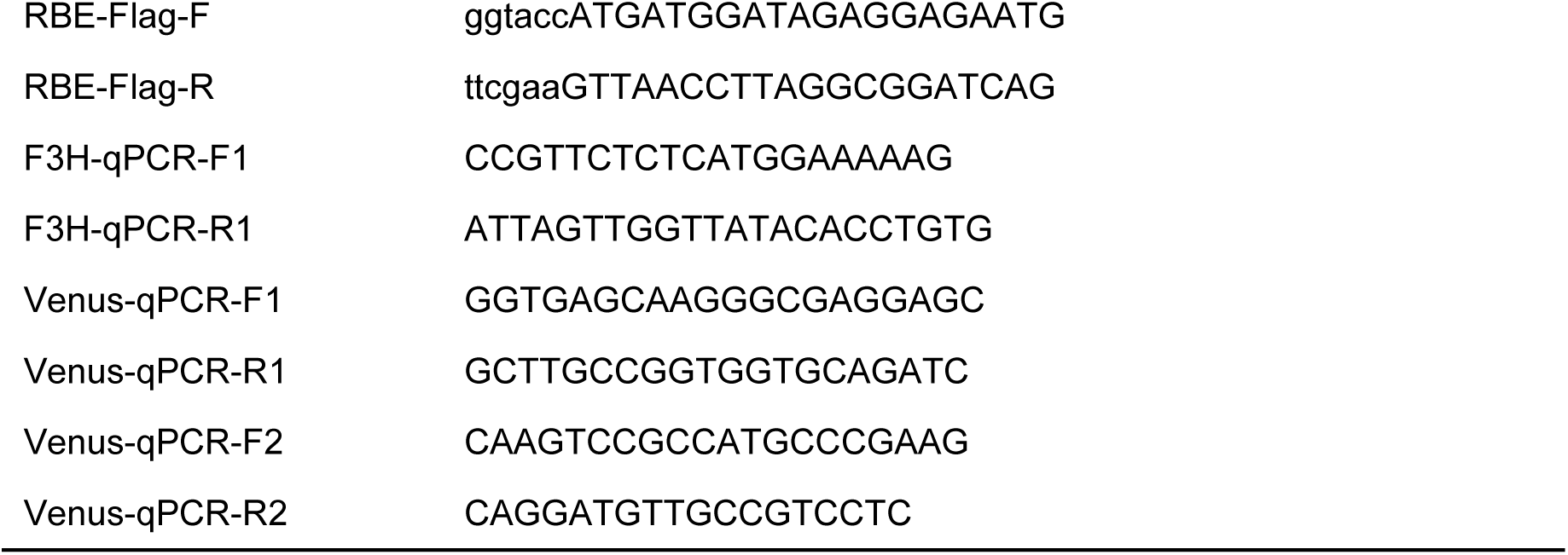
List of primers.

